# PRISME: A MATLAB Toolbox For Large Data-Driven Multimodal Power Benchmarking

**DOI:** 10.64898/2026.01.21.700878

**Authors:** Fabricio Cravo, Alex Fischbach, Hallee Shearer, Matt Rosenblatt, Dustin Scheinost, Stephanie Noble

## Abstract

Low statistical power in neuroimaging often undermines research in the field, leading to missed effects, wasted resources, and reduced reproducibility. Performing power analyses during the study design phase is extremely important, but often prohibitively difficult due to a lack of analytical solutions and high computational costs. We present `PRISME` (Power Resampling Infrastructure for Statistical Method Evaluation), a MATLAB toolbox for neuroimaging power benchmarking. `PRISME` provides a computational framework for empirical power analysis independent of inference methods, enabling large scale power benchmarking and method comparison. The toolbox supports diverse neuroimaging data types, including both voxel-based activation and functional connectivity analyses, with a non-parametric, flexible algorithm and unified data representations. Furthermore, unlike previous empirical power approaches, `PRISME` supports multiple test types, such as association and difference tests with behavioral and clinical measures. Finally, `PRISME`’s 25× speedup from algorithmic optimizations enables larger-scale power benchmarking, including the first power analysis for the ABCD dataset. Overall, `PRISME` is the first method- and data-type-agnostic power benchmarking tool for neuroimaging, providing a single solution for power analysis across diverse study designs.

## 1 Introduction

Answering research questions in neuroscience requires reliably detecting statistically significant patterns called true effects in complex brain imaging data despite substantial noise. [FHW^+^94, MTR^+^12, Hay15, VDHP10, PNPH99, SN18, CB24b, CWY17]. Statistical power, the probability of detecting true effects through methods that compare data to randomness (i.e. statistical inference methods), falls far below the recommended standard of 80% in neuroimaging [NMZS22, SN18, CWY17, BIM^+^13, LAH^+^16], with experiments typically achieving only 8-31% average power [BIM^+^13]. This widespread low power leads to several problems for neuroscience research. First, researchers frequently miss key effects that would validate their hypotheses, resulting in inconclusive findings and inefficient usage of time and resources [BIM^+^13, VKS16, Ell22]. Additionally, low statistical power severely hinders scientific reproducibility, which undermines the ability to distinguish genuine findings from false positives. For example, when an effect has merely 10 % power, only one in ten experimental repetitions is expected to detect it reliably [Coh13]. This may partly explain why replication attempts of neuroimaging studies often fail to reproduce previously reported findings [Ell22, Hen20, MHH18].

A known solution to the low power lies in performing proper a-priori power calculations [SN18, Yeu18]. Through proper power estimation during study design, researchers can determine the sample sizes necessary to reliably detect true effects, identify the magnitude of effects their study is powered to detect, and substantially reduce the probability of missing meaningful results [SN18, SRY^+^24]. Studies that incorporate proper power calculations before data collection demonstrate notably improved research quality, with Mumford [Mum12] showing that the process of conducting power analysis frequently leads researchers to refine their study design and improve experimental tasks. Despite these benefits, the number of studies performing power calculations is concerning. Szucs and Ioannidis [SI20] argued for the importance of power calculation for appropriate allocation of research funding and found that only 3% of studies reported a-priori power calculations in 2017-2018.

Despite the importance of power, the approaches for its estimation in neuroimaging are lacking and do not cover the needs of the field. Some of these are parametric approaches [Mum12, DG02, HPHL07, MN08, DDM^+^16, KMLS09], many of which remain methodological frameworks without implemented software tools [DG02, KMLS09] or have tools that are no longer maintained [Mum12, MN08]. Parametric methods require prior assumptions about the shape of the distribution and the nature of the data to find analytical solutions for calculating power, with different assumptions required for a power calculation for each statistical inference method [Coh13]. If experimental data do not follow these assumptions, the power calculation results will be inaccurate. Furthermore, some popular statistical inference methods in neuroscience, such as the Threshold-Free Cluster Enhancement (TFCE) [SN09] and cluster-size inference [WF95], do not have parametric methods available because they currently lack analytical solutions. In addition to these issues, parametric methods require researchers to specify expected effect sizes a priori, increasing the chances of erroneous power estimation. These limitations increase the likelihood of user errors in power estimation and restrict applicability. Non-parametric methods mitigate these issues by estimating power empirically without requiring distributional assumptions, effect size specification, or analytical solutions for each inference method.

Recognizing these advantages, researchers have developed valuable non-parametric methods for estimating power [NSC20, NMZS22]. Noble et al. [NSC20, NMZS22] proposed a non-parametric subsampling repetition algorithm that uses large datasets to simulate experiments with smaller sample sizes and identifies true effects. Despite its methodological strengths, this approach is not production-ready for widespread use: it lacks computational efficiency, has no standardized framework for method comparison, and is incompatible with diverse data types, datasets, and test types. Even on a computational cluster, it has high computational demands, requiring on average 64 hours without parallelization for just 500 repetitions per study, make it prohibitively expensive for large-scale analyses. Applied to the ABCD dataset with 40 FC studies across 4 different sample sizes (200, 500, 1000, 2000), this would require approximately 1.1 months of computation even with 16 processor cores under highly optimistic computational assumptions (perfect parallelization and equal fitting cost across test types), qualifying the calculation as impractical. Its implementation is hard-coded to Human Connectome Project (HCP) task-based FC studies, lacking support for other datasets, test types beyond the one sample t-test, and voxel activation-based analysis.

Finally, while the previous non-parametric algorithm has been used to compare methods [NSC20, NMZS22], it uses built-in hard-coded inference methods and it has no capability for users to add their own methods easily for analysis. Therefore, there is no standardized framework available for benchmarking multiple statistical inference procedures. Researchers developing new statistical inference methods have no standardized platform to evaluate their methods against established alternatives under identical conditions and would need to code comparisons by themselves with no guarantee of correctness and standardization. Without such a platform, the field lacks an objective basis for determining which statistical methods offer genuine power improvements in detecting neural effects across different experimental contexts.

We present `PRISME`, a MATLAB toolbox for efficient empirical power benchmarking in neuroimaging. PRISME addresses the computational and flexibility limitations of previous approaches through several innovations. First, permutation recycling across methods reduces algorithmic complexity from *O*(Methods × Permutations) to *O*(Methods + Permutations), allowing larger scale and faster power analysis. Second, algorithmic optimizations for widely-used methods like cNBS (134× speedup) and TFCE (34× speedup), combined with support for one-sample, two-sample, and correlation analyses, allow for empirical power analysis at a new scale. We demonstrate this with power estimates across 40 ABCD studies, including brain-behavior correlations and group comparisons. Finally, `PRISME` provides a modular architecture that allows users to add new statistical inference methods easily with a single class script, facilitating the benchmarking of newly developed methods. In the supplementary information, we provide formal mathematical proof of the algorithm’s core concepts relevant for power sub-sampling estimations.

## 2 Results

In this section, we summarize PRISME’s algorithm, computational optimizations, and validation results. First, we describe the subsampling-based power estimation algorithm and the statistical inference methods currently implemented (Section 2.1.2) with the initial GLM fit performed prior (Section 2.1.1). Then, we detail computational optimizations that enabled the large-scale power analysis (Section 2.2). Next, we describe PRISME’s method- and data-agnostic design. Finally, we validate PRISME by replicating previous empirical power estimations for the HCP dataset and extending to 40 ABCD studies and HCP voxel data.

### 2.1 Algorithm

#### 2.1.1 Mass Univariate Regression

`PRISME` performs mass univariate regression by fitting the same statistical model independently to each brain variable (edge or voxel). It uses the multi-stage General Linear Model (GLM) [WF95, NH02, NS20, SN09, ZFB10], generating one t-statistic per variable for the entire subject group.

Let **Y**_*i*_ ∈ ℝ ^1×*n*^ be the data matrix containing measurements for variable numbered *i* (edges or voxels) across *n* subjects, and **X** ∈ ℝ ^*n*×*b*^ the design matrix with *b* model parameters. The design matrix can be a column of ones for one-sample t-tests, a two-column indicator matrix for two-sample t-tests, or continuous measures with an intercept for correlation-based tests. For each variable, we perform the following fit:

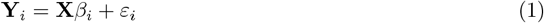

where *β*_*i*_ ∈ ℝ ^*b*^ are the model parameters fit for variable numbered *i*, and ***ε***_*i*_ ∈ ℝ ^1×*n*^ contains the error terms. From each *β*_*i*_ ∈ ℝ ^*b*^, we compute a single t-statistic value. Higher t-statistics indicate stronger evidence against the null hypothesis of no effect. An effect is detected when the null hypothesis has been rejected.

#### 2.1.2 Architecture

The power estimation procedure implemented in `PRISME` is structured around repeated subsampling to evaluate the ability of statistical inference methods to detect effects known to be present in the full dataset (Figure 1). To do so, it utilizes two distinct computational paths: a repeated-sampling loop that evaluates the methods in multiple subsets of the data, and a single estimation step that defines the ground truth effect map. The results of both paths are then combined to provide error rate estimates.

**Figure 1.**
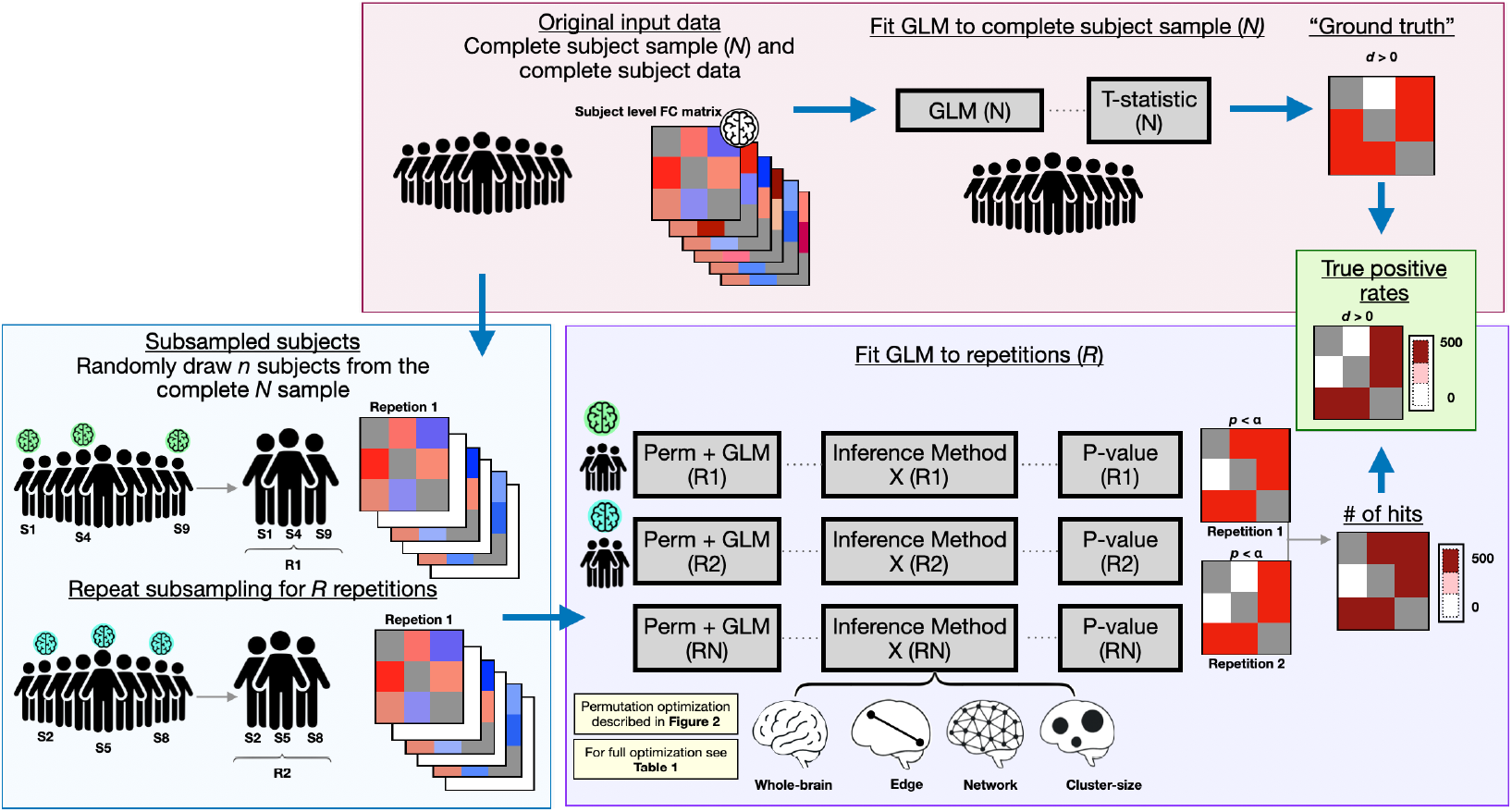
Algorithm workflow for non-parametric power estimation. The workflow comprises two paths for power calculation. **Pink panel**: Complete subject sample path uses all subjects to fit GLM and establish ground truth effect directions based on t-statistic signs. **Blue panel**: Repeated subsampling path randomly draws n subjects from the full dataset across R repetitions. **Purple Panel**: The GLM is fit to both original and permuted data to generate t-statistics, then applies statistical inference methods that produce p-values at different levels (whole-brain, network, edge, cluster-size). Permutations are recycled across all methods for computational efficiency (complexity reduction from O(R×n×M) to O(R×(n+M))). **Green Panel**: Power calculation compares detected effects from each repetition against ground truth, and the statistical power is the proportion of correctly detected effects

In the repeated-sampling path, *n* subjects are randomly sampled without replacement from the larger pool of *N* available subject-level connectivity matrices. The t-statistics are obtained with mass univariate regression in accordance to Section 2.1.1, which are then used by the statistical inference methods described below. The statistical inference methods, in turn, produce p-values for each repetition. For each repetition, p-values are compared to a significance threshold to identify statistically significant effects (i.e., where the null hypothesis is rejected and effects are detected)

#### 2.1.3 Statistical inference methods

`PRISME` evaluates statistical power at three inference levels:

- **Variable level**: Power to detect effects in individual analysis units: FC links (edges) between brain regions for FC data, or individual voxels for activation data.
- **Cluster level**: Power to detect effects in spatially contiguous groups of variables through pooled inference (e.g., using Cluster Size or TFCE methods)
- **Network level**: Power to detect aggregate effects within variables in predefined large-scale functional systems (e.g., default mode, salience networks)
- **Whole-brain level**: Power to detect any significant effect across all variables, i.e. if an effect can be detected in any of the studied variables.

The number of p-values that each repetition produces depends on the inference level: one per variable (variable-level and cluster-level methods), one per network (network-level methods), or a single value (whole-brain inference).

Furthermore, several statistical inference methods representing different levels of brain analysis have currently been implemented for use by `PRISME`. At the individual variable level (single brain connections or voxels), the **Univariate (Parametric)** method calculates p-values by comparing observed t-statistics to the standard t-distribution [WF95, NH02]. **Cluster Size** inference first thresholds the map at a pre-defined statistical threshold, then groups remaining connected variables into clusters and determines significance based on cluster size compared to permutation-generated null clusters [WF95]. **Threshold-Free Cluster Enhancement (TFCE)** combines information from each variable with its neighboring variables and creates spatially-weighted statistics that reflect both local activation strength and spatial extent [SN09]. At the Network level, the **Constrained Network-Based Statistic (cNBS)** pools statistical information from predefined brain networks by averaging t-statistics within each network and comparing these averages against permutation-based null distributions [NS20, ZFB10]. At the whole-brain level, the **Multivariate-cNBS (mv-cNBS)** uses cNBS statistics to form a single vector test statistic; like other multivariate tests, this test reveals a joint effect across brain areas but not at any particular spatial location [NH02]. For methods involving multiple tests, both **False Discovery Rate (FDR)** and **Family-Wise Error Rate (FWER)** corrections are implemented [BH95] according to number of tests at each inference level. FDR controls the expected proportion of false positives among significant results, while FWER controls the probability of making at least one false positive. In total, counting correction methods separately, seven inferential procedures are currently implemented (Parametric-FDR, Parametric-FWER, cNBS-FDR, cNBS-FWER, Cluster Size, TFCE, and mv-cNBS).

#### 2.1.4 Ground truth effects

Separately, ground truth effects are computed using the full sample of *N* subjects by fitting the group-level GLM and computing t-statistics for all variables, and averaged across networks for network-level inference. With the explicit assumption that ground truth effects define true effects [NMZS22], edges and networks are labeled as true effects in the direction of their t-statistic sign. As noted by Turner et al. [TPMB18], ground truth estimated from finite samples may not precisely capture all effect sizes. However, this provides conservative power estimates: including uncertain small effects in ground truth increases detection difficulty, thereby evaluating the capability of inference methods to detect weak effects. For whole-brain level inference, there are no directional effects. If the ground truth effects indicate that any variable has a non-zero t-statistic, then the positives resulting from inference methods at this level are considered true positives.

#### 2.1.5 Ground truth and Sub-sampling Comparison

At the end of the workflow, detected effects from each repetition are compared to ground truth effects. Statistical power for a given brain area or whole-brain test is defined as the proportion of correctly identified effects across all repetitions. This yields power probability estimates for all inference methods for each variables, networks, or whole-brain according to the method’s inference level.

#### 2.1.6 Power Definitions

In this section, we formally describe the equations used for power calculations.

##### Variable and Network Level Power

Let *R* be the number of repetitions. Let *x* be a variable (edge or voxel) or network, and let *t*_GT_(*x*) denote the t-statistic for variable *x* computed from the full sample of *N* subjects For each repetition *r* ∈ {1, …, *R*}, let *D*_*r*_(*x*) ∈ {−1, 0, +1} indicate whether a negative effect, no effect, or positive effect was detected for variable *x*.

Power for variable *x* is defined as:

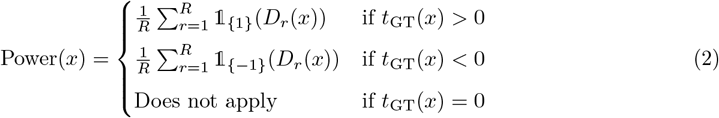

where 𝟙_{*v*}_(*D*_*r*_(*x*)) denotes the indicator function equal to 1 if *D*_*r*_(*x*) = *v* and 0 otherwise, with *v* ∈ {−1, +1}. That is, power measures the proportion of repetitions where the statistical inference method correctly detected an effect in variable *x* in the same direction as the ground truth. Variable *x* is considered to have no effect when *t*_GT_(*x*) = 0. For analysis of `PRISME` applied to variables with no true effect, see Section 7.

##### Whole-Brain Level Power

For whole-brain inference, if ground truth shows any non-zero effect (i.e., ∃*x* : *t*_GT_(*x*)*≠* 0), then power is the proportion of repetitions where any effect was detected. Let *B*_*r*_(*x*) ∈ {0, 1} indicate whether an effect was detected or not, the power at the whole brain level is:

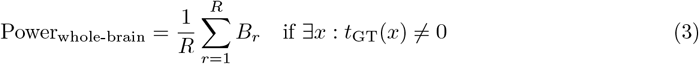

where ∃ is the existence quantifier. Power is undefined if no effects exist in ground truth (null everywhere).

##### Average Power

Let *X* be the set of variables whose variables *x* have a non zero ground truth t-statistic *t*_*GT*_ (*x*) ≠ 0. The average power is:

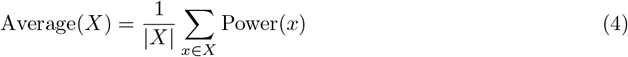

where |*X*| is the number of variables in set *X*.

### 2.2 Computational Optimizations and Performance Analysis

`PRISME` implements several computational optimizations that significantly reduce processing time for non-parametric power estimation. Figure 2 summarizes the computational optimizations and their effects. The key optimizations enabling these performance improvements include:

- **Permutation and GLM fit re-utilization:** Rather than re-fitting the GLM at each permutation for each statistical procedure, `PRISME` performs a single set of permutation-based GLM fits that are then shared across all inference methods. This reduces computational complexity from *O*(Repetitions × Methods × Permutation) to *O*(Repetitions × (Methods + Permutations))
- **Multiple Correction Methods Applied Simultaneously:** When an inference method supports multiple correction procedures (FWER and FDR), uncorrected p-values are computed once, and both corrections can be applied to the same output. This sub-method architecture avoids running the inference method twice, once per correction type.
- **Batch Processing with Check-pointing:** The repetition loop is segmented into configurable batches that automatically save intermediate results after each batch completes. The batch size allows users to control the amount of repetition-related data loaded into RAM for efficiency, with higher batch sizes requiring more RAM. `PRISME` can resume calculations from the saved data to provide resilience against system failures and allow computations to be performed across multiple sections.
- **Permutation Recycling Facilitates C++ Integration:** Statistical inference methods can be among the most computationally intensive components of `PRISME` power calculations, particularly for methods such as TFCE. The class-based architecture enables methods to be prototyped in MATLAB and then replaced with optimized C++ implementations. Since all methods receive identical permutation data from `PRISME`’s recycling component, MATLAB and C++ versions must produce identical outputs when given the same inputs. This direct comparability enables confident validation and replacement of performance-critical code
- **Algorithm Optimizations:** `PRISME` provides faster versions of statistical inference methods. For cNBS, the implementation processes each variable once and assigns it to its network, rather than searching all variables for each network. This reduces the computational complexity from *O*(Networks × Variables) to *O*(Variables). The TFCE algorithm uses an incremental cluster approach that reuses previously computed clusters to compute subsequent threshold clusters. These algorithmic improvements, combined with C++ implementations, provided substantial speedups compared to previous implementations [NS20, SN09] (Table 1).

**Figure 2.**
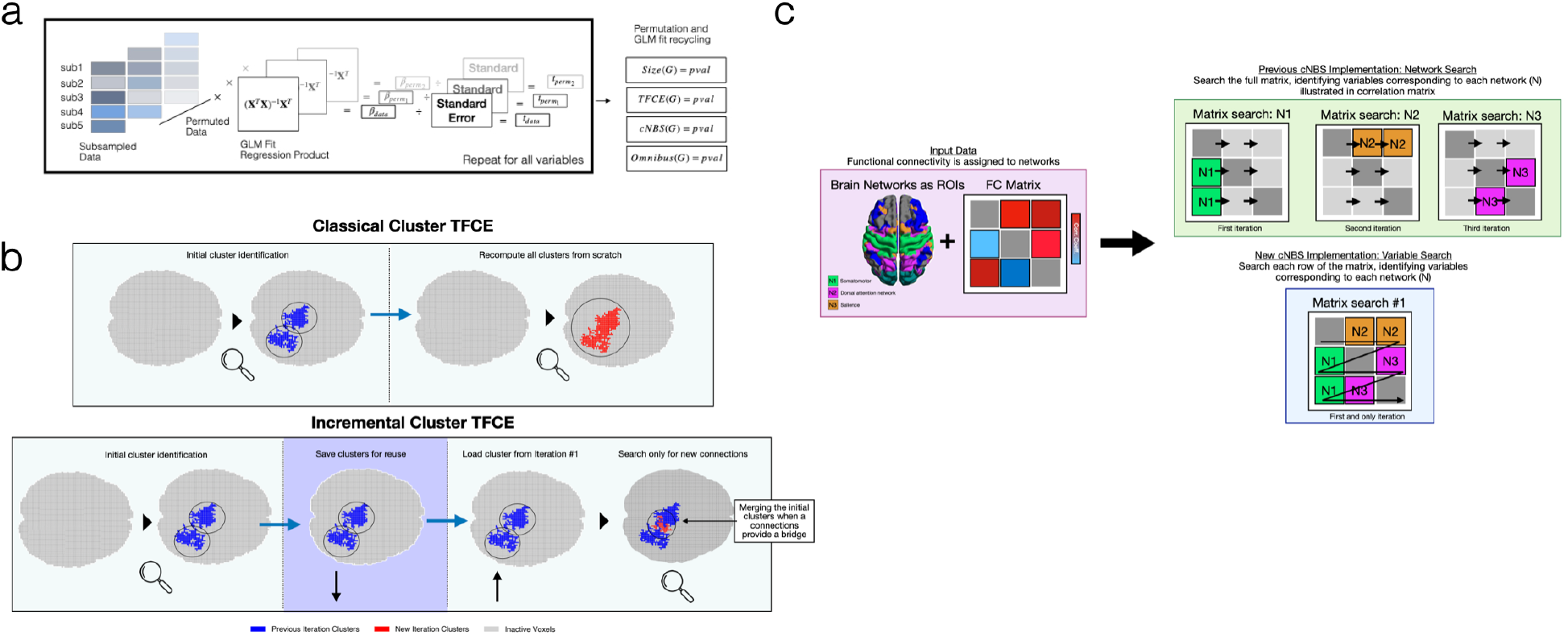
Graphical summary of Computational Optimizations. (a) The GLM fit recycling pipeline. The pipeline shows how GLM fits for both data and permutations of the data are generated and recycled for usage in different methods for computational efficiency. (b) Incremental cluster TFCE implementation. Different from traditional TFCE implementations, `PRISME` computes clusters incrementally for TFCE calculation. (c) cNBS variable level checking. As opposed to the original cNBS implementation that searches for variables within a network for each network, `PRISME`’s implementation loops over all variables once and assigns them to their respective network.

**Table 1:** Performance gains from algorithmic optimizations.

| Computational Improvement | Code Component Speedup Factor |
| --- | --- |
| *Overall Improvement* |  |
| GLM Fit Recycle<br>(5 permutation methods, positive and negative effect direction) | 10× |
| *Method-Specific Computational Improvements* |  |
| Parametric - using same uncorrected p values for FWER + FDR | ~2× |
| Threshold-Free Cluster Enhancement (TFCE) C++ implementation and incremental cluster computation | 34× |
| Cluster Size C++ implementation | 4× |
| cNBS C++ implementation and variable level check and network assignment | 134× |
| **Overall Performance Gain** |  |
| Full HCP dataset (25 studies, 25 cores) | ~25× |

To show `PRISME` computational speed improvements, we performed a computational speed benchmark test. We first conducted a power calculation using the HCP dataset for FC with 35778 edges (Shen 268 atlas [STPC13]) for 5 studies across 5 sample sizes (*n* = 20, 40, 80, 120, 200). Afterwards, we used the reported average time per study reported by Noble et al. [NMZS22] for an estimation of the total time to conduct the same power calculation under similar computing conditions. The estimated time using Noble et al.’s [NMZS22] approach was 20 days, compared to 20 hours with PRISME resulting in a 25× speed up factor (Table 1). Supplementary Method 6.5 contains more details about the benchmark test and computing conditions.

The computational advantages would be greater for other test types. HCP studies primarily use one-sample t-tests requiring the matrix inversion of a single column design matrix, while ABCD studies include two-sample t-tests and correlation analyses with 2-column design matrices that require more computational time to compute the matrix inversion. As PRISME recycles GLM fits across all inference methods rather than recomputing for each method, the efficiency gains for ABCD datasets would exceed the 25× speedup benchmarked on HCP. However, it is impossible for us to compare this as Noble et al.’s was not capable of performing these tests.

### 2.3 Method Agnostic Approach

The `PRISME` toolbox provides the first method-agnostic design within inference levels for neuroimaging power analysis (Figure 3). All that is required is that a method generate p-values from t-statistics at one of the supported inference levels. This method-agnosticism arises from two key design principles. First, method-agnosticism is an inherent consequence of the subsampling approach. By repeatedly subsampling a small number of subjects from the full dataset, the algorithm recreates experimental conditions where any statistical inference method can be applied. The statistical power of each method is then evaluated against ground truth effects estimated from the complete dataset. Second, `PRISME` possesses a modular architecture that facilitates the addition of new methods. Inference methods can be added as new classes that receive t-statistics from upstream code and return p-values.

**Figure 3.**
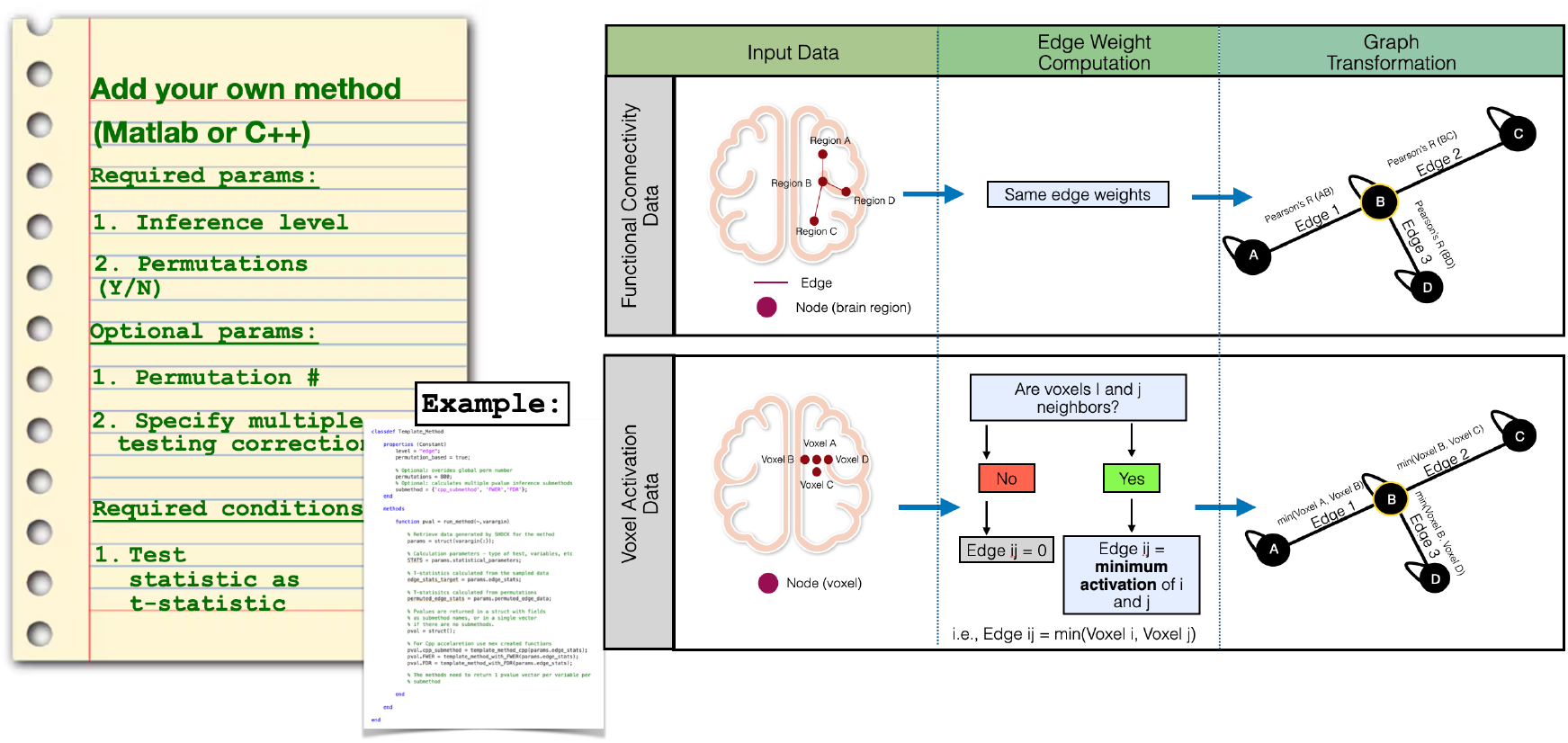
Method- and data-agnostic design of PRISME. *Left* : Method-agnostic interface that allows easy addition of new statistical inference methods. New methods require only specification of inference level (edge/network/whole-brain), permutation requirements, and the core statistical function mapping t-statistics to p-values. *Right* : Data-agnostic method to represent data for methods that require topological information. The figure shows how to process both functional connectivity and voxel activation data through a unified graph transformation. Functional connectivity data map brain regions to graph nodes with correlation strengths as edge weights. Voxel activation data map individual voxels to nodes, with edge weights computed as the minimum activation between spatially adjacent voxels.

### 2.4 Data Agnostic Approach

`PRISME` implements a data-agnostic framework that handles diverse neuroimaging data types through a single pipeline (Figure 3). The data-agnosticism stems from two design principles: flattened array representation and graph-based spatial transformations to abstract statistical inference methods that rely on topological information.

#### 2.4.1 Unified Data Representation

`PRISME` processes both network and image data using flattened arrays, treating FC edges and voxels as the same kind of input variables (Figure 3). The flattened data are able to be directly used without modification by methods that require no spatial information across data types (Parametric, cNBS, and Omnibus).

#### 2.4.2 Graph-Based Spatial Transformation

For inference methods requiring topological relationships (TFCE and Cluster Size), `PRISME` employs a graph-based transformation that converts both data types into an identical graph structure. This transformation allows the same algorithms to operate on either data type by mapping different data types to a common data representation.

The graph construction differs by data type and here we introduce the construction:

##### FC data

Regions of interest become graph nodes, and the FC data between these regions become weighted edges in the graph, with connectivity strengths as edge weights.

##### Voxel activation data

In traditional voxel-based analysis, TFCE and cluster-size methods first threshold the brain at a statistical cutoff (e.g., *t >* 2.5), removing voxels below this threshold, then identify spatially contiguous groups of above-threshold voxels as clusters. A cluster is a set of voxels where each voxel is spatially adjacent to at least one other voxel in the set.

Our graph representation achieves this by creating a graph where voxels become nodes, spatial adjacencies become edges, edge weights equal the minimum activation between connected voxels, and clusters, equal to the FC representation, are connected subgraphs. As an example, consider a case where two spatially adjacent voxels have activations of 3 and 2. In the graph representation, both those voxels would be connected by an edge with the minimum weight of 2. If the threshold is 2.5, this edge falls below the threshold and is excluded from the graph, resulting in the voxels not being connected anymore and not forming a cluster. This ensures the graph-based cluster detection produces identical results to traditional spatial clustering.

Furthermore, `PRISME` precomputes voxel neighborhood relationships during initialization, storing them as a sparse adjacency matrix that represents the graph. This one-time preprocessing enables a fast construction of weighted graphs from flattened activation data using the precomputed adjacency structure.

### 2.5 Power Values and Validation

We validated `PRISME` through two methods. First, we verified that the implementation of statistical inference methods accurately detects known effects using synthetic datasets for each test type (one-sample, two-sample, and association tests). This testing framework confirmed 100% detection accuracy for known synthetic effects and provides method developers with a sanity check for new implementations. Furthermore, this testing framework is implemented as both a standalone script and as a GitHub action.

Second, we replicated and extended the HCP power analysis from Noble et al. [NMZS22] (Figure 4). We analyzed 5 FC task studies using 7 inference methods (Parametric-FDR, Parametric-FWER, cNBS-FDR, cNBS-FWER, Cluster Size, TFCE, and mv-cNBS) across 4 sample sizes (*n* = 250,*n* = 500,*n* = 1000,*n* = 2000) and compared to their original 3 sample sizes (*n* = 40,*n* = 80,*n* = 120).

**Figure 4.**
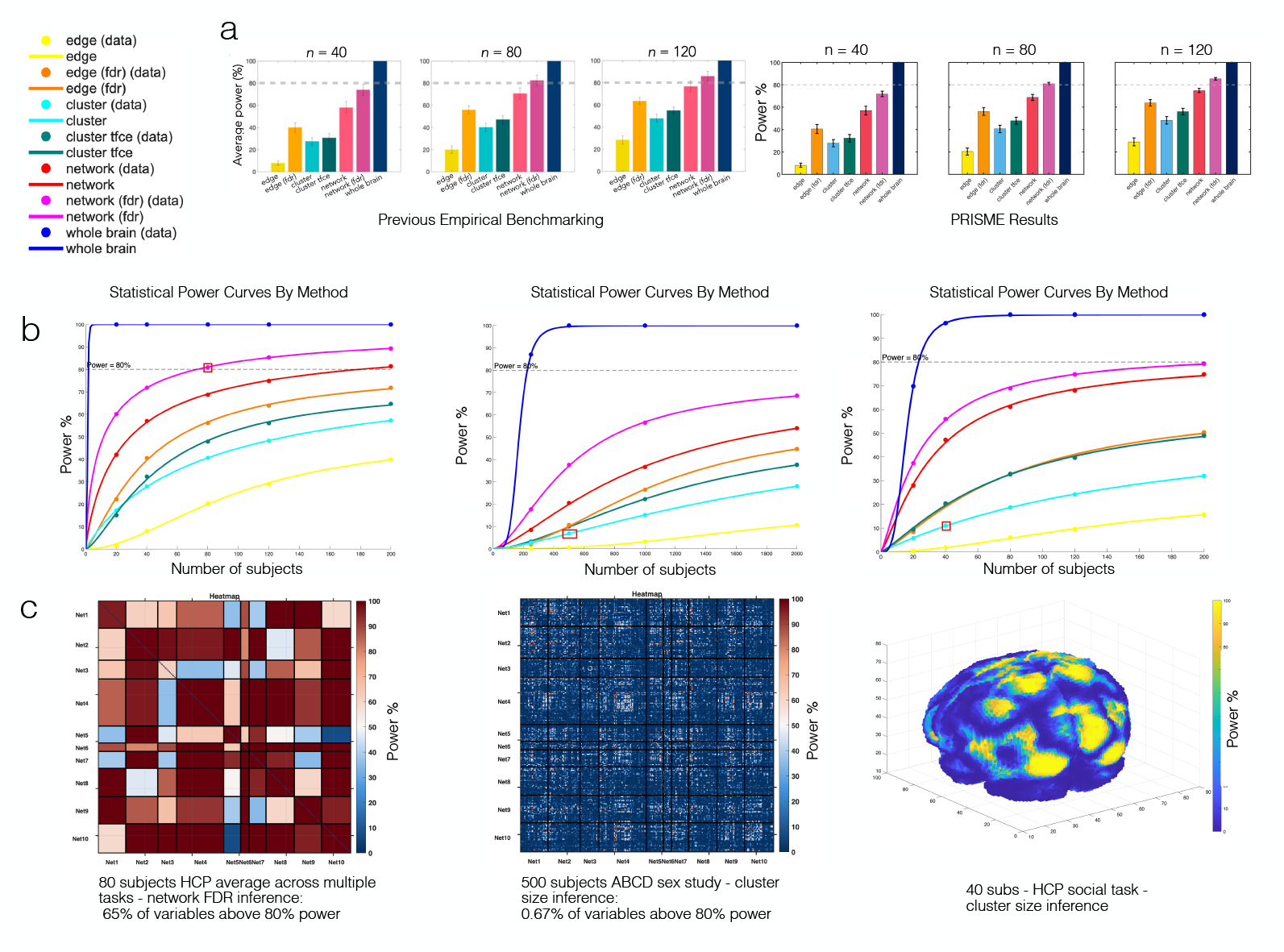
Power analysis results across datasets and methods. (a) Validation of `PRISME` against previous results from Noble et al., showing average statistical power across different sample sizes (n = 40, 80, 120) for seven inference methods. (b) Statistical power curves across three analysis contexts. Left: HCP task-based studies showing power curves fitted to validation data. Middle: ABCD correlation studies (cognitive-FC relationships). Right: Voxel-based analysis showing lower average power but highlighting spatial heterogeneity in effect detection. Red squares indicate the specific sample sizes and method referenced in panel (c). (c) Detailed spatial patterns of specific power analysis.

By visually comparing the histogram plot provided in Figure 4, one can see that the power results from Noble et al. [NMZS22] closely match the results produced by `PRISME`. A direct match is not expected due to the stochastic nature of both approaches for power benchmarking.

For voxel-level analyses, we generated power curves across sample sizes for five HCP tasks (n=20, 40, 80, 120, 200; 100 repetitions). This extends Noble et al. [NSC20], who measured detection rates for different effect sizes at n=20, by determining required sample sizes for 80% power and providing voxel-wise power estimates. We additionally evaluate inference methods not included in their analysis: parametric-FDR, parametric-FWER, cNBS-FDR, cNBS-FWER, and mv-cNBS.

Herein a study is considered a valid and available pairwise comparison between two types of data present in a dataset. For example, FC rest data compared to behavior data as correlation, FC rest data versus task data, and FC rest data versus categorical variables.

Having validated the power benchmarking accuracy of `PRISME`, we demonstrate its computational capability by performing power analyses at a scale previously infeasible: 40 ABCD studies [CCC^+^18, VKC^+^18] (compared to 5 HCP studies in the validation) encompassing both two-sample t-tests (sex differences) and correlation tests (age, cognitive, and behavioral measures). Complete power results for all 40 studies are provided in Supplementary Table 6.2. Figure 4C presents the study with highest observed power: sex differences in resting-state FC, where all methods except cluster size and edge-level parametric FWER exceed 80% power at *n* = 200.

**Table 2:** Power Analysis Results - Biological Sex.

| Method | $n = 250$ | $n = 500$ | $n = 1000$ | $n = 2000$ | a | b | P | $R^2$ |
| --- | --- | --- | --- | --- | --- | --- | --- | --- |
| Network FDR | 17.69 | 37.51 | 56.49 | 68.56 | 511.56 | 1.64 | 75.75 | 1.00 |
| Network FWER | 8.47 | 20.53 | 36.69 | 54.04 | 993.82 | 1.44 | 73.68 | 1.00 |
| TFCE | 3.22 | 10.08 | 22.23 | 37.57 | 1249.15 | 1.66 | 54.71 | 1.00 |
| Omnibus_cNBS | 87.00 | 100.00 | 100.00 | 100.00 | 170.88 | 5.00 | 100.00 | 1.00 |
| Parametric FDR | 2.07 | 10.71 | 26.45 | 44.70 | 1069.16 | 2.02 | 57.22 | 1.00 |
| Parametric FWER | 0.08 | 0.61 | 3.20 | 10.65 | 1855.31 | 2.62 | 19.39 | 1.00 |
| Size | 2.70 | 6.89 | 15.19 | 28.02 | 1996.69 | 1.43 | 55.97 | 1.00 |

Finally, we provide a mathematical proof that validates `PRIME` sub sampling method for power estimation for broad application across multiple data types and statistical inference methods in the supplementary information.

## 3 Methods

### 3.1 Implementation and Availability

`PRISME` is implemented as an open-source MATLAB toolbox (version 2024a or later) distributed under the MIT license via GitHub https://github.com/neuroprismlab/PRISME-Brain-Power-Calculator. The toolbox operates on neuroimaging data in matrix format, with input data structure specifications and data preprocessing detailed in Supplementary Methods. `PRISME` automatically infers the appropriate statistical test type (one-sample, two-sample, or correlation analyses) based on the input data structure. A usage tutorial is available in the Supplementary Methods. Preprocessed neuroimaging data from the Human Connectome Project and Adolescent Brain Cognitive Development Study in PRISME-ready format can be obtained from the authors upon verification of approved data access from the respective repositories.

### 3.2 Statistical Inference Method Implementation

The method agnostic approach of `PRISME` is further strengthened by the simplicity of adding new methods for power benchmarking. The modular architecture allows users to add new inference methods simply by adding a new MATLAB class script with the following attributes:

- The inference level (ex: variable, network, etc …)
- A boolean specifying if permutations are required
- The statistical function converting t-statistics to p-values

This feature facilitates the addition and benchmarking of new methods with minimal code. By handling data processing, permutation generation, and ground truth estimation, `PRISME` allows method developers to focus on statistical innovation while ensuring fair power benchmarking.

## 4 Discussion

`PRISME` is the first general-purpose power benchmarking tool for neuroimaging. Unlike parametric approaches restricted to specific inference methods or previous empirical approaches limited to single data types, PRISME works across multiple methods, data types, and study designs. The algorithm evaluates any inference method by testing its ability to detect effects in subsampled data against ground truth estimated from the more precise full dataset estimation. New methods can be integrated by creating a single new class. PRISME handles both functional connectivity and voxel activation data through a unified framework, supporting one-sample, two-sample, and correlation analyses. For the first time, researchers have a power benchmarking tool for neuroimaging inference methods and data types within a single framework.

Empirical power estimation usually requires extensive computational time and resources, limiting some practical applications. `PRISME` addresses this through three key optimizations. First, permutation recycling shares GLM fits across all inference methods rather than recalculating for each, reducing complexity from *O*(`Methods × Permutations`) to *O*(`Methods`+`Permutations`). Second, some methods are implemented in C++ while maintaining MATLAB compatibility. Third, algorithmic improvements to established methods (cNBS, TFCE) provide additional speedups. Together, these achieve a 25× overall speedup compared to previous empirical approaches, reducing times from weeks to days.

We validated `PRISME` through several analyses. We replicated Noble et al.’s [NMZS22] HCP power analysis, producing power estimates matching previously established results. We used the new computational efficiency to benchmark power on 40 ABCD studies across 4 sample sizes, a tenfold increase in scale compared to previous benchmarking with 5 HCP studies and 3 sample sizes [NMZS22]. Finally, we demonstrated the data-agnostic capability through power analysis of voxel task-activation data. Together, these analyses establish `PRISME` as a general-purpose solution for empirical power estimation in neuroimaging.

The `PRISME` toolbox can extend naturally to other neuroimaging modalities that employ similar statistical approaches. For example, electroencephalogram (EEG) studies utilize functional connectivity analysis [CSA^+^23]. Similarly, functional near-infrared spectroscopy (fNIRS) applies GLM frameworks for task activation and functional connectivity estimation [HZZ^+^21] equally to fMRI. The difference is that fNIRS measures cortical activity near the skull surface with fewer variables than fMRI. These modalities may require only minor data format conversions, or potentially could work with `PRISME` directly without modification. We plan to formalize support for these modalities in future releases.

The ground truth estimation has important limitations. Statistical power is known to increase with the magnitude of the underlying true effect and the sample size [Coh88]. With `PRISME`, we assume the ground truth estimation is the true effect for the entire population. The ground truth effects are estimations from a true effect sampling distribution, whose estimation accuracy increases with the number of subjects in the available dataset [CB24a]. This can lead to errors when the ground truth is not a precise enough estimation of the true underlying effect. These estimation errors directly affect measured power because we measure detection probability relative to the ground truth estimate.

For ground truth effects with incorrectly estimated signs, recent analyses of large neuroimaging datasets reveal that confidence intervals for many effects overlap zero, indicating uncertainty in the true effect sign [SRY^+^24]. This occurs more frequently for low-magnitude effects. `PRISME` partially mitigates this by independently testing for both positive and negative effects. When ground truth incorrectly assigns an effect’s sign, the inference method is more likely to fail to detect the effect across repetitions due to the low magnitude. This results in conservative power estimates, ensuring that actual observed power is more likely to exceed our estimates than fall below them.

`PRISME` addresses a critical gap: the lack of accessible and versatile tools for empirical power analysis by providing the first method- and data-agnostic power benchmarking tool for neuroimaging. With `PRISME`, researchers can perform power calculations for sample size planning, and the framework enables standardized benchmarking of statistical inference methods under equivalent conditions. As neuroimaging continues to grapple with reproducibility challenges, `PRISME` facilitates proper study planning and methodological evaluation to improve research quality. We anticipate that wider adoption of empirical power analysis will contribute to more reliable findings, more efficient resource allocation, and more reproducible neuroscience.

## 5 Acknowledgments

This work was supported by funding from the National Institute of Mental Health (K99 MH130894; R00 MH130894 to S.N).

## 6 Supplementary Methods

### 6.1 Design Matrices and Contrast Matrices

In this section, we detail the design matrices and test coefficients used by `PRISME` for group-level statistical inference following standard approaches [PMN11, Cha20]. We note *n*_*s*_ ∈ ℕ as the number of subjects.

#### One-sample t-test

For testing whether the mean connectivity differs from zero, the design matrix consists of a single column of ones:

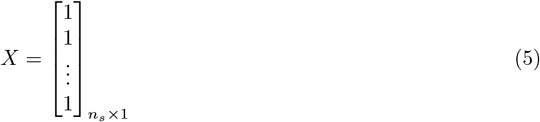

The test coefficients used are: =

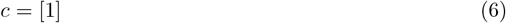

The hypothesis test is:

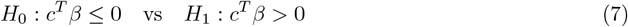

##### Two-sample t-test

For comparing connectivity between two groups, the design matrix uses indicator variables for group membership:

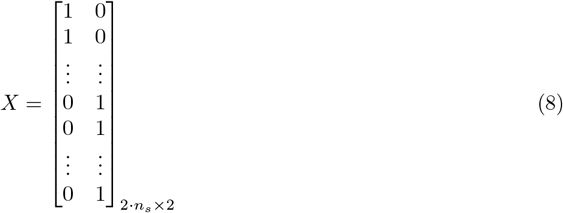

where the first column indicates membership in group 1 and the second column indicates membership in group 2, with *n*_*s*_ = *n*_*g*1_ + *n*_*g*2_, *n*_*g*1_ = *n*_*g*2_ total subjects. The test coefficients for testing if the regressor for group 1 is higher than the regressor for group 2 are:

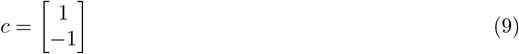

The hypothesis test is:

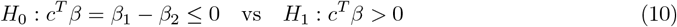

##### Correlation analysis

For testing the relationship between connectivity and a continuous variable (e.g., age, behavioral scores), the design matrix includes the continuous variable and an intercept:

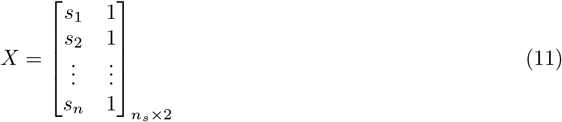

where *s*_*i*_ represents the score or continuous measure for subject *i*, and the second column is the intercept term. The test coefficients used are:

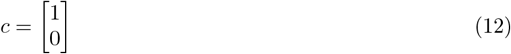

The hypothesis test is:

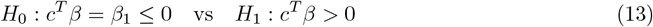

### 6.2 Power Analysis Results - ABCD Dataset

With the new computational improvements from `PRISME`, we performed power analyses on the ABCD dataset [CCC^+^18, VKC^+^18] at a scale previously infeasible by current literature standards. We conducted power calculations across 40 studies using both two-sample t-tests and correlation-based analyses, encompassing 7 inference methods and 4 sample sizes, totaling 1120 empirical power calculations.

The ABCD study outcomes are categorized as follows:

- **Category 1: Demographics & Basic Measures (Studies 1-4)**
  - **Study 1** - `sex`: Biological sex (two-sample t-test)
  - **Study 2** - `pea_wiscv_tss`: WISC-V Total Scale Score (cognitive ability)
  - **Study 3** - `interview_age`: Age at interview
  - **Study 4** - `bmi_z`: Body Mass Index z-score
- **Category 2: CBCL Baseline Measures (Studies 5-14)** Child Behavior Checklist syndrome scales at baseline:
  - **Study 5** - `cbcl_scr_syn_internal_t`: Internalizing problems
  - **Study 6** - `cbcl_scr_syn_external_t`: Externalizing problems
  - **Study 7** - `cbcl_scr_syn_aggressive_t`: Aggressive behavior
  - **Study 8** - `cbcl_scr_syn_rulebreak_t`: Rule-breaking behavior
  - **Study 9** - `cbcl_scr_syn_attention_t`: Attention problems
  - **Study 10** - `cbcl_scr_syn_thought_t`: Thought problems
  - **Study 11** - `cbcl_scr_syn_social_t`: Social problems
  - **Study 12** - `cbcl_scr_syn_somatic_t`: Somatic complaints
  - **Study 13** - `cbcl_scr_syn_withdep_t`: Withdrawn/depressed
  - **Study 14** - `cbcl_scr_syn_anxdep_t`: Anxious/depressed
- **Category 3: UPPS Impulsivity Scale (Studies 15-19)**
  - **Study 15** - `upps_y_ss_lack_of_planning`: Lack of planning
  - **Study 16** - `upps_y_ss_lack_of perseverance`: Lack of perseverance
  - **Study 17** - `upps_y_ss_sensation_seeking`: Sensation seeking
  - **Study 18** - `upps_y_ss_negative_urgency`: Negative urgency
  - **Study 19** - `upps_y_ss_positive_urgency`: Positive urgency
- **Category 4: Substance Use (Study 20)**
  - **Study 20** - `sup_y_ss_sum`: Youth substance use summary score
- **Category 5: CBCL Follow-up 1 (Studies 21-30)** Same CBCL measures at first follow-up visit (FU1):
  - **Studies 21-30**: Mirror studies 5-14 but measured at follow-up
- **Category 6: CBCL Change Scores (Studies 31-40)** Longitudinal change in CBCL measures (delta = FU1 - baseline):
  - **Studies 31-40**: Change scores for the same CBCL domains

Tables 2–41 contain the results for this analysis for all the studies. For all power calculations, the following power curve for the average power across all variables was fit to the experimental data:

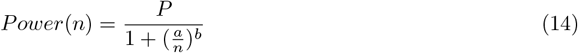

where *n* ∈?ℝ is the number of of subjects and *a, b, P* ∈ ℝ are constants fit to data.

The power analysis results (Tables 2–41) show low average statistical power across all calculations. Aside from whole-brain level inference, the biological sex differences analysis (Table **??**) produced the highest power, but none of the methods reached the commonly recommended 80% threshold. Because the ABCD dataset consists primarily of children, the generally low power in the association tests may reflect the small magnitude of categorical differences in this population. Power would likely be higher in a sample with more variation across subjects.

**Table 3:** Power Analysis Results - WISC-V Total Scale Score.

| Method | $n = 250$ | $n = 500$ | $n = 1000$ | $n = 2000$ | a | b | P | $R^2$ |
| --- | --- | --- | --- | --- | --- | --- | --- | --- |
| Network FDR | 0.73 | 1.27 | 2.98 | 6.89 | 16510.36 | 1.23 | 100.00 | 1.00 |
| Network FWER | 0.56 | 0.87 | 2.00 | 3.96 | 42946.01 | 1.04 | 99.89 | 1.00 |
| TFCE | 0.02 | 0.13 | 0.82 | 4.41 | 3046.63 | 2.75 | 18.49 | 1.00 |
| Omnibus_cNBS | 0.00 | 1.00 | 0.00 | 1.00 | 99959.90 | 0.67 | 12.35 | 0.21 |
| Parametric FDR | 0.01 | 0.05 | 0.50 | 4.65 | 3342.18 | 3.42 | 31.57 | 1.00 |
| Parametric FWER | 0.00 | 0.00 | 0.02 | 0.17 | 15432.52 | 2.85 | 57.29 | 1.00 |
| Size | 0.15 | 0.48 | 1.48 | 3.98 | 3520.44 | 1.73 | 14.59 | 1.00 |

**Table 4:** Power Analysis Results - Age at Interview.

| Method | $n = 250$ | $n = 500$ | $n = 1000$ | $n = 2000$ | a | b | P | $R^2$ |
| --- | --- | --- | --- | --- | --- | --- | --- | --- |
| Network FDR | 0.49 | 0.62 | 1.64 | 6.45 | 8103.05 | 1.91 | 100.00 | 0.99 |
| Network FWER | 0.36 | 0.40 | 1.09 | 3.42 | 16879.06 | 1.57 | 99.99 | 0.99 |
| TFCE | 0.02 | 0.13 | 1.07 | 4.87 | 1826.83 | 3.23 | 8.51 | 1.00 |
| Omnibus_cNBS | 0.00 | 0.00 | 1.00 | 24.00 | 2472.56 | 5.00 | 93.31 | 1.00 |
| Parametric FDR | 0.00 | 0.04 | 0.75 | 5.38 | 1682.84 | 4.34 | 7.92 | 1.00 |
| Parametric FWER | 0.00 | 0.00 | 0.03 | 0.21 | 13534.20 | 2.88 | 51.43 | 1.00 |
| Size | 0.16 | 0.49 | 1.60 | 4.31 | 2998.72 | 1.81 | 13.27 | 1.00 |

**Table 5:** Power Analysis Results - BMI Z-Score.

| Method | $n = 250$ | $n = 500$ | $n = 1000$ | $n = 2000$ | a | b | P | $R^2$ |
| --- | --- | --- | --- | --- | --- | --- | --- | --- |
| Network FDR | 1.75 | 3.40 | 9.51 | 19.29 | 2256.82 | 1.55 | 42.59 | 1.00 |
| Network FWER | 1.24 | 2.36 | 5.64 | 10.64 | 2774.62 | 1.32 | 27.07 | 1.00 |
| TFCE | 0.12 | 1.08 | 4.64 | 12.91 | 1702.55 | 2.43 | 21.62 | 1.00 |
| Omnibus_cNBS | 0.00 | 4.00 | 27.00 | 89.00 | 1250.13 | 4.37 | 100.00 | 1.00 |
| Parametric FDR | 0.06 | 0.81 | 5.06 | 16.46 | 1582.81 | 2.95 | 24.71 | 1.00 |
| Parametric FWER | 0.01 | 0.06 | 0.39 | 1.73 | 2221.74 | 2.81 | 4.05 | 1.00 |
| Size | 0.54 | 1.69 | 4.28 | 9.87 | 3516.52 | 1.51 | 33.03 | 1.00 |

**Table 6:**
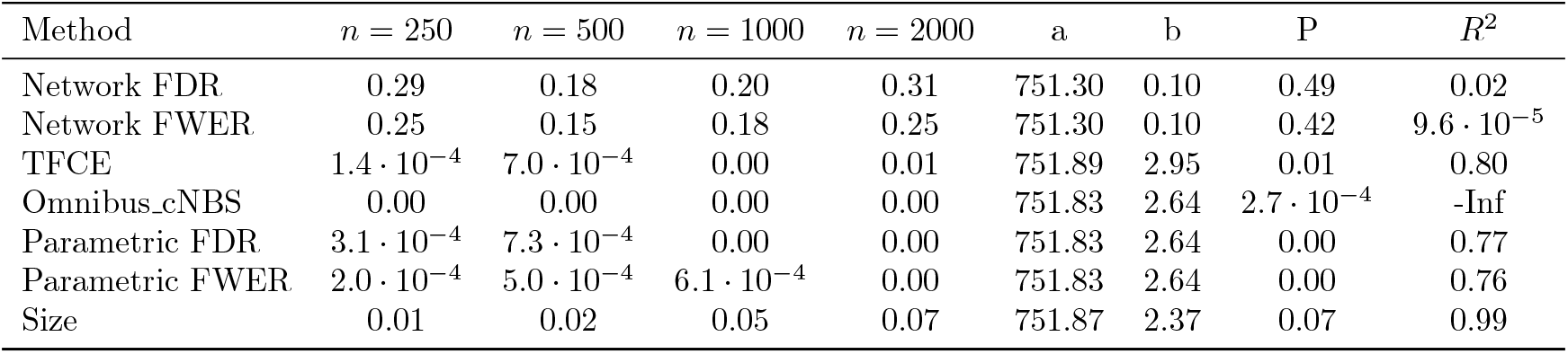
Power Analysis Results - CBCL Internalizing Problems (Baseline)

**Table 7:** Power Analysis Results - CBCL Externalizing Problems (Baseline)

| Method | $n = 250$ | $n = 500$ | $n = 1000$ | $n = 2000$ | a | b | P | $R^2$ |
| --- | --- | --- | --- | --- | --- | --- | --- | --- |
| Network FDR | 0.22 | 0.36 | 0.44 | 0.93 | 99888.08 | 0.78 | 19.90 | 0.95 |
| Network FWER | 0.22 | 0.35 | 0.42 | 0.76 | 99810.71 | 0.65 | 10.32 | 0.96 |
| TFCE | 0.00 | 0.00 | 0.01 | 0.01 | 751.66 | 1.92 | 0.02 | 0.77 |
| Omnibus.cNBS | 1.00 | 0.00 | 0.00 | 0.00 | 1.00 | 0.10 | 0.37 | -0.02 |
| Parametric FDR | 0.00 | 0.00 | 0.00 | 0.01 | 751.41 | 0.69 | 0.01 | 0.33 |
| Parametric FWER | $6.4 \cdot 10^{-4}$ | $5.3 \cdot 10^{-4}$ | 0.00 | 0.00 | 751.83 | 2.63 | 0.00 | 0.63 |
| Size | 0.03 | 0.02 | 0.07 | 0.18 | 90012.61 | 1.32 | 27.33 | 0.97 |

**Table 8:** Power Analysis Results - CBCL Aggressive Behavior (Baseline)

| Method | $n = 250$ | $n = 500$ | $n = 1000$ | $n = 2000$ | a | b | P | $R^2$ |
| --- | --- | --- | --- | --- | --- | --- | --- | --- |
| Network FDR | 0.22 | 0.27 | 0.24 | 0.64 | 99901.16 | 0.69 | 9.41 | 0.77 |
| Network FWER | 0.15 | 0.25 | 0.20 | 0.49 | 99787.33 | 0.64 | 6.08 | 0.78 |
| TFCE | $8.1 \cdot 10^{-4}$ | 0.00 | 0.01 | 0.02 | 751.90 | 3.10 | 0.02 | 0.92 |
| Omnibus.cNBS | 0.00 | 0.00 | 0.00 | 0.00 | 751.83 | 2.64 | $2.7 \cdot 10^{-4}$ | -Inf |
| Parametric FDR | $7.0 \cdot 10^{-4}$ | 0.00 | 0.00 | 0.01 | 751.82 | 2.63 | 0.01 | 0.71 |
| Parametric FWER | $4.5 \cdot 10^{-4}$ | $6.4 \cdot 10^{-4}$ | $7.0 \cdot 10^{-4}$ | 0.00 | 751.83 | 2.63 | 0.00 | 0.65 |
| Size | 0.02 | 0.03 | 0.06 | 0.18 | 87579.16 | 1.33 | 27.16 | 0.99 |

**Table 9:** Power Analysis Results - CBCL Rule-Breaking Behavior (Baseline)

| Method | $n = 250$ | $n = 500$ | $n = 1000$ | $n = 2000$ | a | b | P | $R^2$ |
| --- | --- | --- | --- | --- | --- | --- | --- | --- |
| Network FDR | 0.47 | 0.44 | 0.58 | 1.53 | 99998.93 | 0.89 | 48.53 | 0.87 |
| Network FWER | 0.36 | 0.31 | 0.45 | 1.20 | 99959.82 | 0.94 | 46.19 | 0.88 |
| TFCE | 0.01 | 0.01 | 0.03 | 0.17 | 24883.05 | 2.32 | 58.12 | 1.00 |
| Omnibus.cNBS | 1.00 | 0.00 | 0.00 | 1.00 | 751.31 | 0.10 | 1.00 | -0.00 |
| Parametric FDR | 0.00 | 0.00 | 0.01 | 0.05 | 9427.62 | 2.08 | 1.40 | 0.99 |
| Parametric FWER | 0.00 | $7.8 \cdot 10^{-4}$ | 0.00 | 0.01 | 751.79 | 2.48 | 0.01 | 0.75 |
| Size | 0.05 | 0.06 | 0.19 | 0.64 | 39391.86 | 1.68 | 96.51 | 1.00 |

**Table 10:**
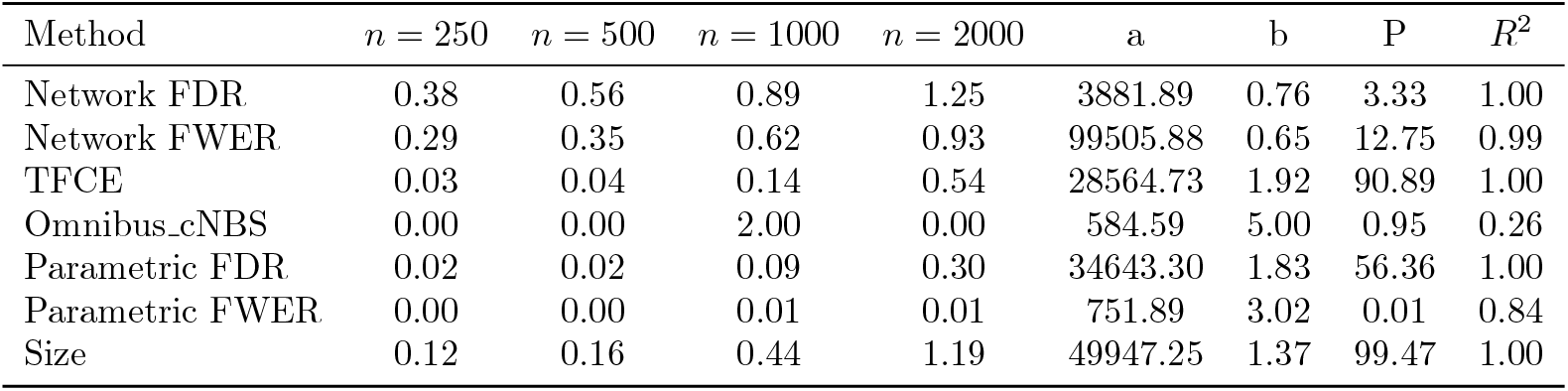
Power Analysis Results - CBCL Attention Problems (Baseline)

**Table 11:**
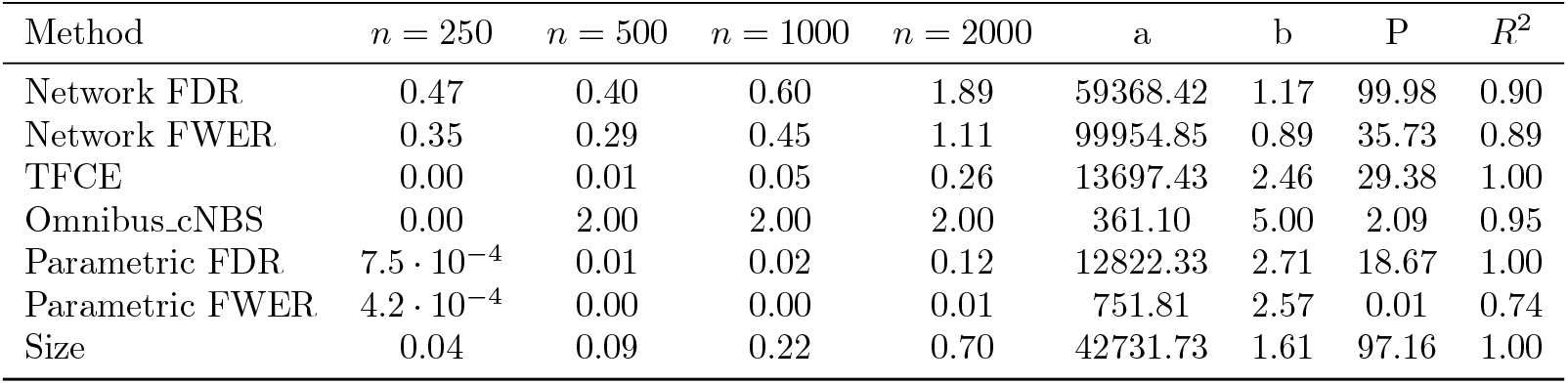
Power Analysis Results - CBCL Thought Problems (Baseline)

| Method | $n = 250$ | $n = 500$ | $n = 1000$ | $n = 2000$ | a | b | P | $R^2$ |
| --- | --- | --- | --- | --- | --- | --- | --- | --- |
| Network FDR | 0.47 | 0.40 | 0.60 | 1.89 | 59368.42 | 1.17 | 99.98 | 0.90 |
| Network FWER | 0.35 | 0.29 | 0.45 | 1.11 | 99954.85 | 0.89 | 35.73 | 0.89 |
| TFCE | 0.00 | 0.01 | 0.05 | 0.26 | 13697.43 | 2.46 | 29.38 | 1.00 |
| Omnibus.cNBS | 0.00 | 2.00 | 2.00 | 2.00 | 361.10 | 5.00 | 2.09 | 0.95 |
| Parametric FDR | $7.5 \cdot 10^{-4}$ | 0.01 | 0.02 | 0.12 | 12822.33 | 2.71 | 18.67 | 1.00 |
| Parametric FWER | $4.2 \cdot 10^{-4}$ | 0.00 | 0.00 | 0.01 | 751.81 | 2.57 | 0.01 | 0.74 |
| Size | 0.04 | 0.09 | 0.22 | 0.70 | 42731.73 | 1.61 | 97.16 | 1.00 |

**Table 12:** Power Analysis Results - CBCL Social Problems (Baseline)

| Method | $n = 250$ | $n = 500$ | $n = 1000$ | $n = 2000$ | a | b | P | $R^2$ |
| --- | --- | --- | --- | --- | --- | --- | --- | --- |
| Network FDR | 0.11 | 0.27 | 0.45 | 0.44 | 411.75 | 2.65 | 0.46 | 0.98 |
| Network FWER | 0.09 | 0.25 | 0.33 | 0.35 | 354.17 | 2.94 | 0.35 | 1.00 |
| TFCE | 0.02 | 0.01 | 0.01 | 0.08 | 4374.81 | 3.31 | 1.16 | 0.91 |
| Omnibus.cNBS | 0.00 | 0.00 | 0.00 | 0.00 | 751.83 | 2.64 | $2.7 \cdot 10^{-4}$ | -Inf |
| Parametric FDR | 0.01 | 0.00 | 0.00 | 0.03 | 751.55 | 1.45 | 0.03 | 0.31 |
| Parametric FWER | 0.00 | 0.00 | 0.00 | 0.00 | 751.63 | 1.75 | 0.00 | 0.63 |
| Size | 0.07 | 0.08 | 0.10 | 0.36 | 98361.82 | 1.34 | 65.42 | 0.91 |

**Table 13:**
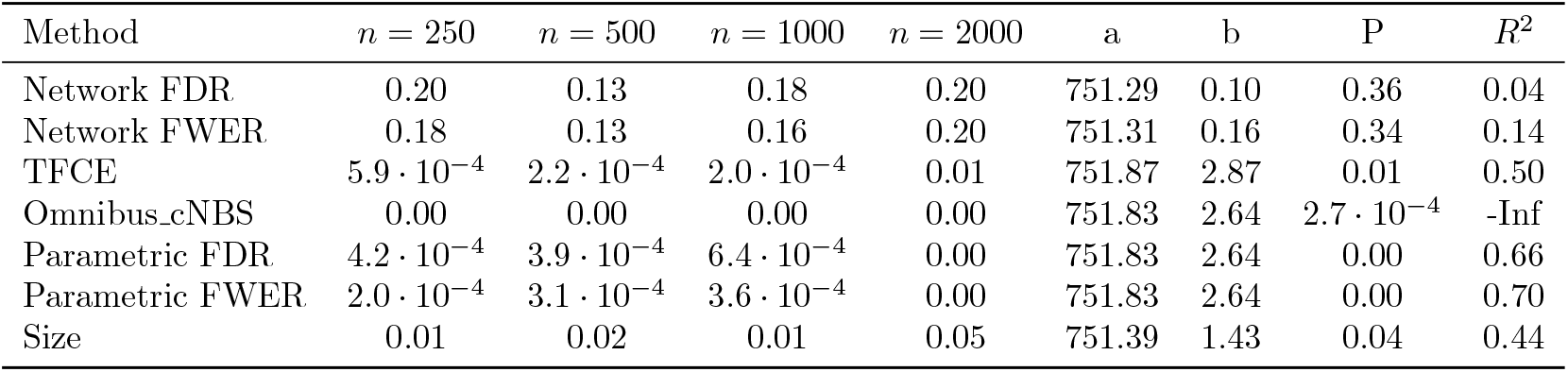
Power Analysis Results - CBCL Somatic Complaints (Baseline)

| Method | $n = 250$ | $n = 500$ | $n = 1000$ | $n = 2000$ | a | b | P | $R^2$ |
| --- | --- | --- | --- | --- | --- | --- | --- | --- |
| Network FDR | 0.20 | 0.13 | 0.18 | 0.20 | 751.29 | 0.10 | 0.36 | 0.04 |
| Network FWER | 0.18 | 0.13 | 0.16 | 0.20 | 751.31 | 0.16 | 0.34 | 0.14 |
| TFCE | $5.9 \cdot 10^{-4}$ | $2.2 \cdot 10^{-4}$ | $2.0 \cdot 10^{-4}$ | 0.01 | 751.87 | 2.87 | 0.01 | 0.50 |
| Omnibus.cNBS | 0.00 | 0.00 | 0.00 | 0.00 | 751.83 | 2.64 | $2.7 \cdot 10^{-4}$ | -Inf |
| Parametric FDR | $4.2 \cdot 10^{-4}$ | $3.9 \cdot 10^{-4}$ | $6.4 \cdot 10^{-4}$ | 0.00 | 751.83 | 2.64 | 0.00 | 0.66 |
| Parametric FWER | $2.0 \cdot 10^{-4}$ | $3.1 \cdot 10^{-4}$ | $3.6 \cdot 10^{-4}$ | 0.00 | 751.83 | 2.64 | 0.00 | 0.70 |
| Size | 0.01 | 0.02 | 0.01 | 0.05 | 751.39 | 1.43 | 0.04 | 0.44 |

**Table 14:** Power Analysis Results - CBCL Withdrawn/Depressed (Baseline)

| Method | $n = 250$ | $n = 500$ | $n = 1000$ | $n = 2000$ | a | b | P | $R^2$ |
| --- | --- | --- | --- | --- | --- | --- | --- | --- |
| Network FDR | 0.24 | 0.29 | 0.29 | 0.24 | 751.30 | 0.10 | 0.53 | -0.14 |
| Network FWER | 0.20 | 0.22 | 0.24 | 0.24 | 751.31 | 0.16 | 0.45 | 0.89 |
| TFCE | $10.0 \cdot 10^{-4}$ | 0.01 | 0.01 | 0.01 | 751.64 | 1.82 | 0.01 | 0.98 |
| Omnibus.cNBS | 0.00 | 0.00 | 1.00 | 0.00 | 584.59 | 5.00 | 0.47 | 0.26 |
| Parametric FDR | $3.9 \cdot 10^{-4}$ | 0.00 | 0.00 | 0.01 | 751.72 | 2.18 | 0.01 | 0.99 |
| Parametric FWER | $2.0 \cdot 10^{-4}$ | $5.6 \cdot 10^{-4}$ | 0.00 | 0.00 | 751.83 | 2.64 | 0.00 | 0.95 |
| Size | 0.02 | 0.03 | 0.04 | 0.10 | 97055.00 | 1.01 | 5.20 | 0.98 |

**Table 15:** Power Analysis Results - CBCL Anxious/Depressed (Baseline)

| Method | $n = 250$ | $n = 500$ | $n = 1000$ | $n = 2000$ | a | b | P | $R^2$ |
| --- | --- | --- | --- | --- | --- | --- | --- | --- |
| Network FDR | 0.27 | 0.11 | 0.47 | 0.20 | 751.31 | 0.16 | 0.53 | 0.01 |
| Network FWER | 0.18 | 0.07 | 0.25 | 0.18 | 751.34 | 0.31 | 0.35 | 0.10 |
| TFCE | 0.07 | 0.03 | 0.04 | 0.01 | 751.28 | 0.10 | 0.08 | -0.12 |
| Omnibus.cNBS | 0.00 | 0.00 | 2.00 | 0.00 | 584.59 | 5.00 | 0.95 | 0.26 |
| Parametric FDR | 0.06 | 0.01 | 0.02 | 0.00 | 751.33 | 0.10 | 0.05 | -0.07 |
| Parametric FWER | 0.00 | 0.00 | 0.00 | 0.00 | 751.32 | 0.26 | 0.00 | -0.67 |
| Size | 0.14 | 0.08 | 0.15 | 0.12 | 751.29 | 0.10 | 0.25 | 0.00 |

**Table 16:** Power Analysis Results - UPPS Lack of Planning.

| Method | $n = 250$ | $n = 500$ | $n = 1000$ | $n = 2000$ | a | b | P | $R^2$ |
| --- | --- | --- | --- | --- | --- | --- | --- | --- |
| Network FDR | 0.24 | 0.38 | 0.16 | 0.95 | 99947.53 | 1.05 | 53.36 | 0.67 |
| Network FWER | 0.18 | 0.20 | 0.15 | 0.51 | 99867.09 | 0.77 | 9.86 | 0.67 |
| TFCE | $6.4 \cdot 10^{-4}$ | 0.00 | 0.00 | 0.02 | 751.88 | 2.96 | 0.02 | 0.62 |
| Omnibus.cNBS | 0.00 | 0.00 | 0.00 | 0.00 | 751.83 | 2.64 | $2.7 \cdot 10^{-4}$ | -Inf |
| Parametric FDR | $7.0 \cdot 10^{-4}$ | $8.7 \cdot 10^{-4}$ | 0.00 | 0.01 | 751.79 | 2.48 | 0.00 | 0.68 |
| Parametric FWER | $3.4 \cdot 10^{-4}$ | $4.8 \cdot 10^{-4}$ | $7.3 \cdot 10^{-4}$ | 0.00 | 751.83 | 2.64 | 0.00 | 0.74 |
| Size | 0.00 | 0.02 | 0.06 | 0.17 | 3864.15 | 1.75 | 0.71 | 1.00 |

**Table 17:** Power Analysis Results - UPPS Lack of Perseverance.

| Method | $n = 250$ | $n = 500$ | $n = 1000$ | $n = 2000$ | a | b | P | $R^2$ |
| --- | --- | --- | --- | --- | --- | --- | --- | --- |
| Network FDR | 0.13 | 0.09 | 0.04 | 0.13 | 751.29 | 0.10 | 0.19 | -0.04 |
| Network FWER | 0.11 | 0.07 | 0.04 | 0.09 | 751.29 | 0.10 | 0.15 | -0.10 |
| TFCE | 0.00 | $5.7 \cdot 10^{-4}$ | 0.00 | 0.04 | 751.87 | 3.12 | 0.03 | 0.59 |
| Omnibus.cNBS | 0.00 | 0.00 | 0.00 | 0.00 | 751.83 | 2.64 | $2.7 \cdot 10^{-4}$ | -Inf |
| Parametric FDR | $5.6 \cdot 10^{-4}$ | $7.5 \cdot 10^{-4}$ | 0.00 | 0.02 | 751.88 | 2.99 | 0.02 | 0.56 |
| Parametric FWER | $3.6 \cdot 10^{-4}$ | $4.8 \cdot 10^{-4}$ | $6.7 \cdot 10^{-4}$ | 0.00 | 751.83 | 2.64 | 0.00 | 0.65 |
| Size | 0.02 | 0.02 | 0.06 | 0.23 | 43949.68 | 1.88 | 76.75 | 0.99 |

**Table 18:** Power Analysis Results - UPPS Sensation Seeking.

| Method | $n = 250$ | $n = 500$ | $n = 1000$ | $n = 2000$ | a | b | P | $R^2$ |
| --- | --- | --- | --- | --- | --- | --- | --- | --- |
| Network FDR | 0.24 | 0.09 | 0.22 | 0.47 | 99842.71 | 0.72 | 7.89 | 0.63 |
| Network FWER | 0.16 | 0.05 | 0.18 | 0.45 | 99494.06 | 1.06 | 27.71 | 0.81 |
| TFCE | 0.00 | 0.01 | 0.00 | 0.02 | 751.62 | 1.69 | 0.01 | 0.50 |
| Omnibus.cNBS | 0.00 | 0.00 | 0.00 | 0.00 | 751.83 | 2.64 | $2.7 \cdot 10^{-4}$ | -Inf |
| Parametric FDR | 0.00 | 0.00 | 0.00 | 0.01 | 751.55 | 1.33 | 0.01 | 0.52 |
| Parametric FWER | $5.9 \cdot 10^{-4}$ | $5.6 \cdot 10^{-4}$ | $7.8 \cdot 10^{-4}$ | 0.00 | 751.83 | 2.63 | 0.00 | 0.55 |
| Size | 0.02 | 0.03 | 0.03 | 0.14 | 92754.94 | 1.62 | 67.93 | 0.92 |

**Table 19:** Power Analysis Results - UPPS Negative Urgency.

| Method | $n = 250$ | $n = 500$ | $n = 1000$ | $n = 2000$ | a | b | P | $R^2$ |
| --- | --- | --- | --- | --- | --- | --- | --- | --- |
| Network FDR | 0.36 | 0.29 | 0.85 | 2.42 | 27441.35 | 1.41 | 99.97 | 0.98 |
| Network FWER | 0.27 | 0.24 | 0.60 | 1.53 | 59584.31 | 1.23 | 99.90 | 0.97 |
| TFCE | $5.8 \cdot 10^{-4}$ | 0.00 | 0.01 | 0.10 | 5669.61 | 3.27 | 3.04 | 1.00 |
| Omnibus.cNBS | 0.00 | 0.00 | 0.00 | 0.00 | 751.83 | 2.64 | $2.7 \cdot 10^{-4}$ | -Inf |
| Parametric FDR | $4.2 \cdot 10^{-4}$ | $8.1 \cdot 10^{-4}$ | 0.00 | 0.03 | 751.90 | 3.09 | 0.02 | 0.60 |
| Parametric FWER | $3.6 \cdot 10^{-4}$ | $4.2 \cdot 10^{-4}$ | 0.00 | 0.00 | 751.83 | 2.64 | 0.00 | 0.80 |
| Size | 0.01 | 0.04 | 0.12 | 0.41 | 32694.32 | 1.80 | 62.41 | 1.00 |

**Table 20:**
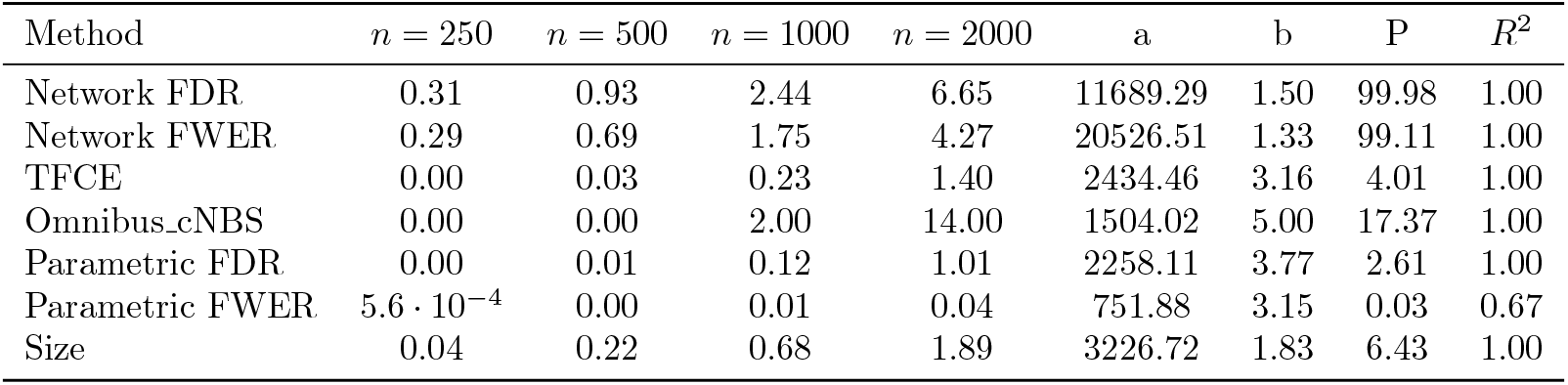
Power Analysis Results - UPPS Positive Urgency.

| Method | $n = 250$ | $n = 500$ | $n = 1000$ | $n = 2000$ | a | b | P | $R^2$ |
| --- | --- | --- | --- | --- | --- | --- | --- | --- |
| Network FDR | 0.31 | 0.93 | 2.44 | 6.65 | 11689.29 | 1.50 | 99.98 | 1.00 |
| Network FWER | 0.29 | 0.69 | 1.75 | 4.27 | 20526.51 | 1.33 | 99.11 | 1.00 |
| TFCE | 0.00 | 0.03 | 0.23 | 1.40 | 2434.46 | 3.16 | 4.01 | 1.00 |
| Omnibus.cNBS | 0.00 | 0.00 | 2.00 | 14.00 | 1504.02 | 5.00 | 17.37 | 1.00 |
| Parametric FDR | 0.00 | 0.01 | 0.12 | 1.01 | 2258.11 | 3.77 | 2.61 | 1.00 |
| Parametric FWER | $5.6 \cdot 10^{-4}$ | 0.00 | 0.01 | 0.04 | 751.88 | 3.15 | 0.03 | 0.67 |
| Size | 0.04 | 0.22 | 0.68 | 1.89 | 3226.72 | 1.83 | 6.43 | 1.00 |

**Table 21:** Power Analysis Results - Substance Use Summary.

| Method | $n = 250$ | $n = 500$ | $n = 1000$ | $n = 2000$ | a | b | P | $R^2$ |
| --- | --- | --- | --- | --- | --- | --- | --- | --- |
| Network FDR | 0.60 | 1.07 | 2.67 | 7.20 | 11940.55 | 1.43 | 100.00 | 1.00 |
| Network FWER | 0.40 | 0.71 | 1.93 | 4.58 | 20390.77 | 1.30 | 99.23 | 1.00 |
| TFCE | 0.01 | 0.10 | 0.36 | 2.08 | 9211.45 | 2.51 | 97.83 | 1.00 |
| Omnibus.cNBS | 0.00 | 0.00 | 1.00 | 12.00 | 1775.32 | 5.00 | 18.61 | 1.00 |
| Parametric FDR | 0.01 | 0.06 | 0.18 | 1.70 | 7034.65 | 3.17 | 93.79 | 1.00 |
| Parametric FWER | 0.00 | 0.00 | 0.01 | 0.06 | 7925.84 | 2.62 | 2.27 | 1.00 |
| Size | 0.13 | 0.33 | 0.91 | 2.38 | 9819.25 | 1.48 | 27.54 | 1.00 |

**Table 22:** Power Analysis Results - CBCL Internalizing Problems (Follow-up)

| Method | $n = 250$ | $n = 500$ | $n = 1000$ | $n = 2000$ | a | b | P | $R^2$ |
| --- | --- | --- | --- | --- | --- | --- | --- | --- |
| Network FDR | 0.24 | 0.11 | 0.33 | 0.36 | 99329.50 | 0.42 | 2.28 | 0.52 |
| Network FWER | 0.20 | 0.09 | 0.31 | 0.33 | 98549.57 | 0.46 | 2.33 | 0.54 |
| TFCE | 0.00 | 0.00 | 0.00 | 0.01 | 751.79 | 2.51 | 0.01 | 0.53 |
| Omnibus.cNBS | 0.00 | 0.00 | 0.00 | 0.00 | 751.83 | 2.64 | $2.7 \cdot 10^{-4}$ | -Inf |
| Parametric FDR | $8.9 \cdot 10^{-4}$ | 0.00 | $9.5 \cdot 10^{-4}$ | 0.00 | 751.62 | 1.69 | 0.00 | 0.60 |
| Parametric FWER | $3.4 \cdot 10^{-4}$ | $4.8 \cdot 10^{-4}$ | $5.9 \cdot 10^{-4}$ | 0.00 | 751.83 | 2.64 | 0.00 | 0.67 |
| Size | 0.02 | 0.01 | 0.03 | 0.13 | 11930.66 | 2.15 | 6.20 | 0.97 |

**Table 23:** Power Analysis Results - CBCL Externalizing Problems (Follow-up)

| Method | $n = 250$ | $n = 500$ | $n = 1000$ | $n = 2000$ | a | b | P | $R^2$ |
| --- | --- | --- | --- | --- | --- | --- | --- | --- |
| Network FDR | 0.44 | 0.56 | 0.67 | 1.04 | 99885.37 | 0.48 | 7.56 | 0.96 |
| Network FWER | 0.33 | 0.47 | 0.51 | 0.91 | 99898.40 | 0.55 | 8.31 | 0.92 |
| TFCE | $5.0 \cdot 10^{-4}$ | 0.00 | 0.01 | 0.07 | 2221.43 | 3.19 | 0.17 | 1.00 |
| Omnibus.cNBS | 0.00 | 0.00 | 0.00 | 1.00 | 4998.84 | 5.00 | 98.42 | 1.00 |
| Parametric FDR | $7.5 \cdot 10^{-4}$ | $8.7 \cdot 10^{-4}$ | 0.00 | 0.03 | 751.88 | 3.04 | 0.02 | 0.57 |
| Parametric FWER | $5.0 \cdot 10^{-4}$ | $5.6 \cdot 10^{-4}$ | $8.4 \cdot 10^{-4}$ | 0.00 | 751.83 | 2.64 | 0.00 | 0.66 |
| Size | 0.01 | 0.04 | 0.10 | 0.37 | 35152.10 | 1.84 | 73.44 | 1.00 |

**Table 24:**
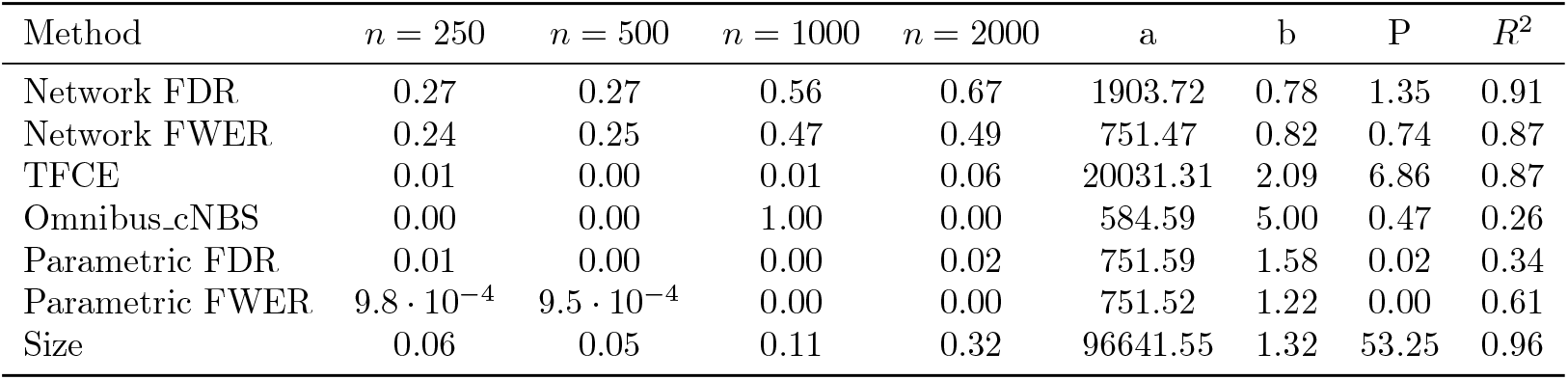
Power Analysis Results - CBCL Aggressive Behavior (Follow-up)

| Method | $n = 250$ | $n = 500$ | $n = 1000$ | $n = 2000$ | a | b | P | $R^2$ |
| --- | --- | --- | --- | --- | --- | --- | --- | --- |
| Network FDR | 0.27 | 0.27 | 0.56 | 0.67 | 1903.72 | 0.78 | 1.35 | 0.91 |
| Network FWER | 0.24 | 0.25 | 0.47 | 0.49 | 751.47 | 0.82 | 0.74 | 0.87 |
| TFCE | 0.01 | 0.00 | 0.01 | 0.06 | 20031.31 | 2.09 | 6.86 | 0.87 |
| Omnibus.cNBS | 0.00 | 0.00 | 1.00 | 0.00 | 584.59 | 5.00 | 0.47 | 0.26 |
| Parametric FDR | 0.01 | 0.00 | 0.00 | 0.02 | 751.59 | 1.58 | 0.02 | 0.34 |
| Parametric FWER | $9.8 \cdot 10^{-4}$ | $9.5 \cdot 10^{-4}$ | 0.00 | 0.00 | 751.52 | 1.22 | 0.00 | 0.61 |
| Size | 0.06 | 0.05 | 0.11 | 0.32 | 96641.55 | 1.32 | 53.25 | 0.96 |

**Table 25:** Power Analysis Results - CBCL Rule-Breaking Behavior (Follow-up)

| Method | $n = 250$ | $n = 500$ | $n = 1000$ | $n = 2000$ | a | b | P | $R^2$ |
| --- | --- | --- | --- | --- | --- | --- | --- | --- |
| Network FDR | 0.24 | 0.71 | 1.09 | 3.36 | 22050.47 | 1.40 | 100.00 | 0.98 |
| Network FWER | 0.24 | 0.55 | 0.85 | 2.11 | 61750.02 | 1.12 | 100.00 | 0.99 |
| TFCE | 0.01 | 0.01 | 0.03 | 0.41 | 7470.64 | 3.72 | 56.15 | 1.00 |
| Omnibus.cNBS | 0.00 | 0.00 | 0.00 | 1.00 | 4998.84 | 5.00 | 98.42 | 1.00 |
| Parametric FDR | 0.01 | 0.00 | 0.01 | 0.19 | 5301.97 | 3.91 | 8.72 | 1.00 |
| Parametric FWER | $7.0 \cdot 10^{-4}$ | $9.5 \cdot 10^{-4}$ | 0.00 | 0.01 | 751.88 | 2.93 | 0.01 | 0.70 |
| Size | 0.06 | 0.12 | 0.25 | 0.97 | 25785.46 | 1.80 | 97.05 | 0.99 |

**Table 26:**
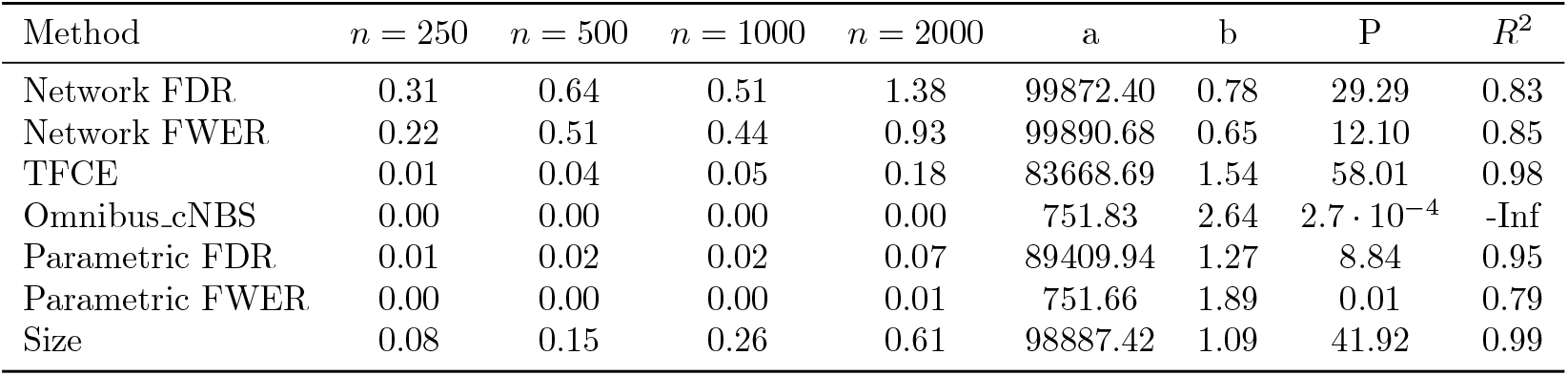
Power Analysis Results - CBCL Attention Problems (Follow-up)

**Table 27:** Power Analysis Results - CBCL Thought Problems (Follow-up)

| Method | $n = 250$ | $n = 500$ | $n = 1000$ | $n = 2000$ | a | b | P | $R^2$ |
| --- | --- | --- | --- | --- | --- | --- | --- | --- |
| Network FDR | 0.38 | 0.33 | 0.56 | 1.18 | 99958.90 | 0.81 | 28.13 | 0.91 |
| Network FWER | 0.31 | 0.31 | 0.40 | 0.78 | 99917.43 | 0.62 | 8.99 | 0.87 |
| TFCE | 0.00 | 0.00 | 0.01 | 0.08 | 4425.63 | 3.83 | 1.80 | 1.00 |
| Omnibus.cNBS | 0.00 | 0.00 | 0.00 | 0.00 | 751.83 | 2.64 | $2.7 \cdot 10^{-4}$ | -Inf |
| Parametric FDR | $7.3 \cdot 10^{-4}$ | 0.00 | 0.00 | 0.04 | 751.83 | 2.89 | 0.03 | 0.57 |
| Parametric FWER | $4.2 \cdot 10^{-4}$ | $8.1 \cdot 10^{-4}$ | 0.00 | 0.00 | 751.79 | 2.50 | 0.00 | 0.77 |
| Size | 0.03 | 0.04 | 0.07 | 0.32 | 36539.29 | 1.94 | 91.01 | 0.98 |

**Table 28:** Power Analysis Results - CBCL Social Problems (Follow-up)

| Method | $n = 250$ | $n = 500$ | $n = 1000$ | $n = 2000$ | a | b | P | $R^2$ |
| --- | --- | --- | --- | --- | --- | --- | --- | --- |
| Network FDR | 0.22 | 0.44 | 0.44 | 0.67 | 29379.55 | 0.52 | 3.33 | 0.90 |
| Network FWER | 0.20 | 0.38 | 0.42 | 0.62 | 3657.34 | 0.64 | 1.51 | 0.94 |
| TFCE | 0.00 | 0.01 | 0.01 | 0.05 | 751.62 | 2.34 | 0.04 | 0.71 |
| Omnibus.cNBS | 1.00 | 0.00 | 0.00 | 0.00 | 1.00 | 0.10 | 0.37 | -0.02 |
| Parametric FDR | 0.00 | 0.00 | 0.00 | 0.02 | 751.76 | 2.37 | 0.01 | 0.63 |
| Parametric FWER | $4.2 \cdot 10^{-4}$ | $9.5 \cdot 10^{-4}$ | 0.00 | 0.00 | 751.82 | 2.63 | 0.00 | 0.72 |
| Size | 0.02 | 0.06 | 0.12 | 0.29 | 81155.52 | 1.22 | 26.48 | 1.00 |

**Table 29:** Power Analysis Results - CBCL Somatic Complaints (Follow-up)

| Method | $n = 250$ | $n = 500$ | $n = 1000$ | $n = 2000$ | a | b | P | $R^2$ |
| --- | --- | --- | --- | --- | --- | --- | --- | --- |
| Network FDR | 0.24 | 0.25 | 0.27 | 0.44 | 99572.96 | 0.38 | 2.21 | 0.81 |
| Network FWER | 0.24 | 0.20 | 0.24 | 0.33 | 751.36 | 0.36 | 0.51 | 0.54 |
| TFCE | 0.00 | $8.9 \cdot 10^{-4}$ | 0.00 | 0.01 | 751.61 | 1.63 | 0.01 | 0.74 |
| Omnibus.cNBS | 0.00 | 0.00 | 0.00 | 0.00 | 751.83 | 2.64 | $2.7 \cdot 10^{-4}$ | -Inf |
| Parametric FDR | $3.1 \cdot 10^{-4}$ | $5.3 \cdot 10^{-4}$ | 0.00 | 0.00 | 751.83 | 2.64 | 0.00 | 0.97 |
| Parametric FWER | $2.5 \cdot 10^{-4}$ | $2.5 \cdot 10^{-4}$ | $5.0 \cdot 10^{-4}$ | $6.4 \cdot 10^{-4}$ | 751.83 | 2.64 | $7.9 \cdot 10^{-4}$ | 0.50 |
| Size | 0.02 | 0.01 | 0.03 | 0.05 | 751.63 | 1.30 | 0.06 | 0.73 |

**Table 30:** Power Analysis Results - CBCL Withdrawn/Depressed (Follow-up)

| Method | $n = 250$ | $n = 500$ | $n = 1000$ | $n = 2000$ | a | b | P | $R^2$ |
| --- | --- | --- | --- | --- | --- | --- | --- | --- |
| Network FDR | 0.07 | 0.13 | 0.20 | 0.13 | 751.39 | 0.52 | 0.27 | 0.34 |
| Network FWER | 0.07 | 0.11 | 0.18 | 0.13 | 751.40 | 0.57 | 0.25 | 0.45 |
| TFCE | $5.9 \cdot 10^{-4}$ | $3.6 \cdot 10^{-4}$ | 0.00 | 0.01 | 751.91 | 3.06 | 0.01 | 0.64 |
| Omnibus.cNBS | 0.00 | 0.00 | 0.00 | 0.00 | 751.83 | 2.64 | $2.7 \cdot 10^{-4}$ | -Inf |
| Parametric FDR | $4.8 \cdot 10^{-4}$ | $6.4 \cdot 10^{-4}$ | 0.00 | 0.00 | 751.80 | 2.51 | 0.00 | 0.89 |
| Parametric FWER | $2.5 \cdot 10^{-4}$ | $5.6 \cdot 10^{-4}$ | $8.9 \cdot 10^{-4}$ | 0.00 | 751.83 | 2.64 | 0.00 | 0.89 |
| Size | 0.02 | 0.01 | 0.03 | 0.11 | 74646.72 | 1.71 | 52.76 | 0.97 |

**Table 31:** Power Analysis Results - CBCL Anxious/Depressed (Follow-up)

| Method | $n = 250$ | $n = 500$ | $n = 1000$ | $n = 2000$ | a | b | P | $R^2$ |
| --- | --- | --- | --- | --- | --- | --- | --- | --- |
| Network FDR | 0.13 | 0.15 | 0.27 | 0.62 | 99607.13 | 1.01 | 32.06 | 0.98 |
| Network FWER | 0.09 | 0.15 | 0.24 | 0.60 | 99428.36 | 1.12 | 46.69 | 0.98 |
| TFCE | 0.00 | 0.00 | $6.7 \cdot 10^{-4}$ | 0.01 | 751.66 | 1.88 | 0.01 | 0.47 |
| Omnibus.cNBS | 0.00 | 0.00 | 0.00 | 0.00 | 751.83 | 2.64 | $2.7 \cdot 10^{-4}$ | -Inf |
| Parametric FDR | 0.00 | $8.4 \cdot 10^{-4}$ | $6.1 \cdot 10^{-4}$ | 0.01 | 751.79 | 2.47 | 0.01 | 0.52 |
| Parametric FWER | $5.0 \cdot 10^{-4}$ | $5.9 \cdot 10^{-4}$ | $4.2 \cdot 10^{-4}$ | 0.00 | 751.83 | 2.63 | 0.00 | 0.44 |
| Size | 0.02 | 0.03 | 0.02 | 0.12 | 96457.80 | 1.43 | 28.61 | 0.87 |

**Table 32:**
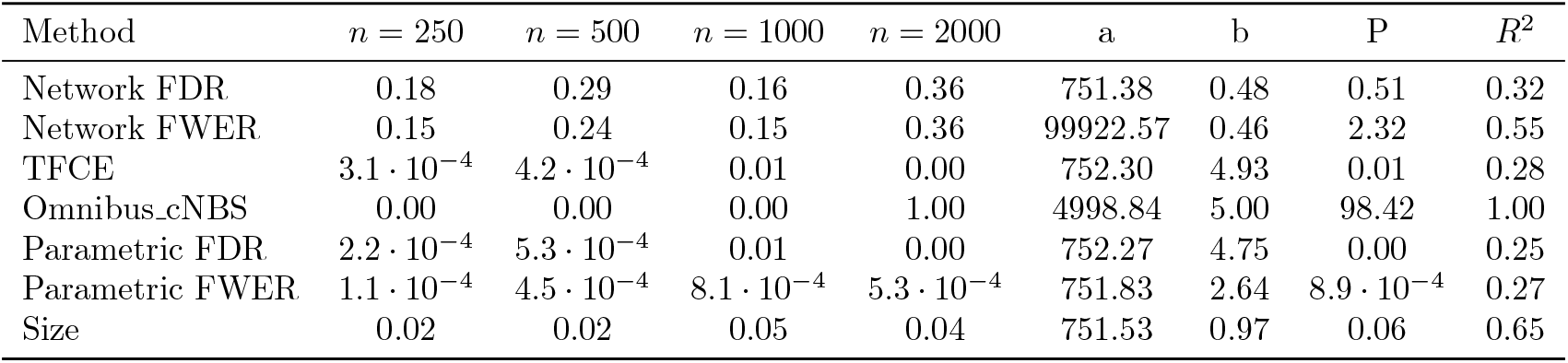
Power Analysis Results - CBCL Internalizing Problems (Change)

| Method | $n = 250$ | $n = 500$ | $n = 1000$ | $n = 2000$ | a | b | P | $R^2$ |
| --- | --- | --- | --- | --- | --- | --- | --- | --- |
| Network FDR | 0.18 | 0.29 | 0.16 | 0.36 | 751.38 | 0.48 | 0.51 | 0.32 |
| Network FWER | 0.15 | 0.24 | 0.15 | 0.36 | 99922.57 | 0.46 | 2.32 | 0.55 |
| TFCE | $3.1 \cdot 10^{-4}$ | $4.2 \cdot 10^{-4}$ | 0.01 | 0.00 | 752.30 | 4.93 | 0.01 | 0.28 |
| Omnibus.cNBS | 0.00 | 0.00 | 0.00 | 1.00 | 4998.84 | 5.00 | 98.42 | 1.00 |
| Parametric FDR | $2.2 \cdot 10^{-4}$ | $5.3 \cdot 10^{-4}$ | 0.01 | 0.00 | 752.27 | 4.75 | 0.00 | 0.25 |
| Parametric FWER | $1.1 \cdot 10^{-4}$ | $4.5 \cdot 10^{-4}$ | $8.1 \cdot 10^{-4}$ | $5.3 \cdot 10^{-4}$ | 751.83 | 2.64 | $8.9 \cdot 10^{-4}$ | 0.27 |
| Size | 0.02 | 0.02 | 0.05 | 0.04 | 751.53 | 0.97 | 0.06 | 0.65 |

**Table 33:**
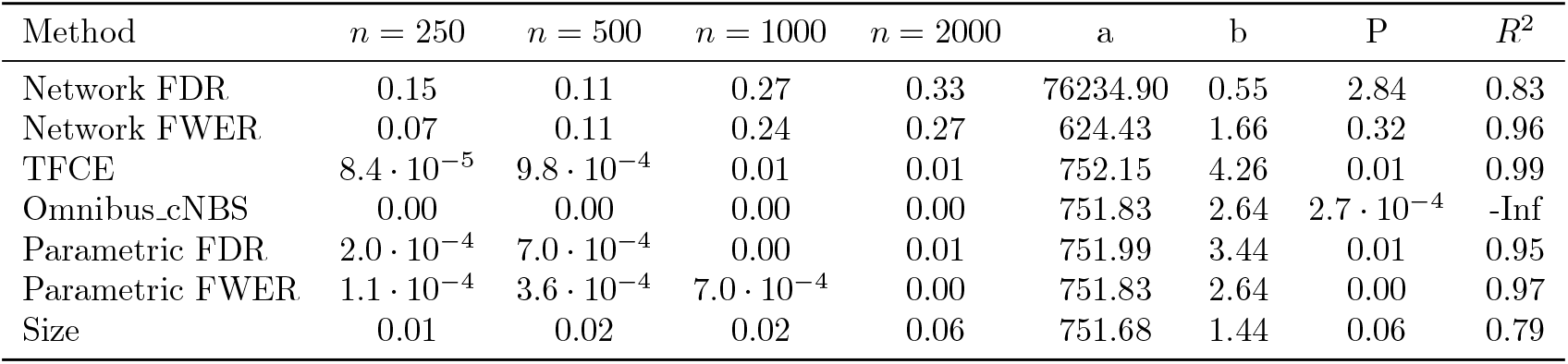
Power Analysis Results - CBCL Externalizing Problems (Change)

**Table 34:** Power Analysis Results - CBCL Aggressive Behavior (Change)

| Method | $n = 250$ | $n = 500$ | $n = 1000$ | $n = 2000$ | a | b | P | $R^2$ |
| --- | --- | --- | --- | --- | --- | --- | --- | --- |
| Network FDR | 0.25 | 0.27 | 0.24 | 0.16 | 751.30 | 0.10 | 0.46 | -0.41 |
| Network FWER | 0.18 | 0.16 | 0.22 | 0.16 | 751.29 | 0.10 | 0.36 | -0.10 |
| TFCE | 0.00 | 0.00 | 0.00 | 0.00 | 751.33 | 0.30 | 0.00 | -0.65 |
| Omnibus.cNBS | 0.00 | 0.00 | 0.00 | 0.00 | 751.83 | 2.64 | $2.7 \cdot 10^{-4}$ | -Inf |
| Parametric FDR | 0.00 | 0.00 | 0.00 | 0.00 | 751.34 | 0.34 | 0.00 | -0.20 |
| Parametric FWER | $7.5 \cdot 10^{-4}$ | $7.3 \cdot 10^{-4}$ | $9.5 \cdot 10^{-4}$ | $7.3 \cdot 10^{-4}$ | 751.82 | 2.63 | 0.00 | -22.86 |
| Size | 0.04 | 0.03 | 0.03 | 0.05 | 751.34 | 0.31 | 0.08 | 0.32 |

**Table 35:** Power Analysis Results - CBCL Rule-Breaking Behavior (Change)

| Method | $n = 250$ | $n = 500$ | $n = 1000$ | $n = 2000$ | a | b | P | $R^2$ |
| --- | --- | --- | --- | --- | --- | --- | --- | --- |
| Network FDR | 0.22 | 0.16 | 0.24 | 0.09 | 751.29 | 0.10 | 0.35 | -0.16 |
| Network FWER | 0.13 | 0.15 | 0.16 | 0.09 | 751.29 | 0.10 | 0.26 | -0.18 |
| TFCE | $7.0 \cdot 10^{-4}$ | $2.8 \cdot 10^{-4}$ | 0.01 | $3.1 \cdot 10^{-4}$ | 752.28 | 4.79 | 0.00 | 0.14 |
| Omnibus.cNBS | 1.00 | 0.00 | 1.00 | 0.00 | 1.00 | 0.10 | 0.75 | -0.02 |
| Parametric FDR | $3.6 \cdot 10^{-4}$ | $4.8 \cdot 10^{-4}$ | 0.00 | $8.9 \cdot 10^{-4}$ | 751.83 | 2.64 | 0.00 | 0.23 |
| Parametric FWER | $1.7 \cdot 10^{-4}$ | $3.4 \cdot 10^{-4}$ | $4.8 \cdot 10^{-4}$ | $5.6 \cdot 10^{-4}$ | 751.83 | 2.64 | $7.4 \cdot 10^{-4}$ | 0.37 |
| Size | 0.02 | 0.01 | 0.04 | 0.01 | 751.30 | 0.11 | 0.04 | -0.00 |

**Table 36:** Power Analysis Results - CBCL Attention Problems (Change)

| Method | $n = 250$ | $n = 500$ | $n = 1000$ | $n = 2000$ | a | b | P | $R^2$ |
| --- | --- | --- | --- | --- | --- | --- | --- | --- |
| Network FDR | 0.20 | 0.11 | 0.29 | 0.13 | 751.29 | 0.10 | 0.36 | -0.02 |
| Network FWER | 0.18 | 0.11 | 0.27 | 0.13 | 751.29 | 0.10 | 0.35 | -0.01 |
| TFCE | 0.07 | 0.02 | 0.01 | 0.01 | 751.23 | 0.10 | 0.05 | -0.08 |
| Omnibus.cNBS | 0.00 | 1.00 | 0.00 | 0.00 | 751.29 | 0.10 | 0.49 | -0.01 |
| Parametric FDR | 0.08 | 0.01 | 0.01 | 0.00 | 751.33 | 0.10 | 0.05 | -0.05 |
| Parametric FWER | 0.00 | 0.00 | 0.00 | 0.00 | 751.82 | 2.60 | 0.00 | -8.73 |
| Size | 0.12 | 0.10 | 0.09 | 0.07 | 751.29 | 0.10 | 0.19 | -0.46 |

**Table 37:** Power Analysis Results - CBCL Thought Problems (Change)

| Method | $n = 250$ | $n = 500$ | $n = 1000$ | $n = 2000$ | a | b | P | $R^2$ |
| --- | --- | --- | --- | --- | --- | --- | --- | --- |
| Network FDR | 0.18 | 0.15 | 0.15 | 0.24 | 751.33 | 0.27 | 0.36 | 0.24 |
| Network FWER | 0.16 | 0.15 | 0.15 | 0.20 | 751.31 | 0.19 | 0.33 | 0.30 |
| TFCE | 0.01 | 0.00 | 0.01 | 0.01 | 751.30 | 0.15 | 0.01 | -0.02 |
| Omnibus.cNBS | 0.00 | 0.00 | 2.00 | 0.00 | 584.59 | 5.00 | 0.95 | 0.26 |
| Parametric FDR | 0.00 | 0.00 | 0.00 | 0.00 | 751.35 | 0.41 | 0.00 | 0.03 |
| Parametric FWER | $8.9 \cdot 10^{-4}$ | $6.4 \cdot 10^{-4}$ | $7.5 \cdot 10^{-4}$ | $8.9 \cdot 10^{-4}$ | 751.82 | 2.63 | 0.00 | -18.49 |
| Size | 0.06 | 0.02 | 0.04 | 0.07 | 751.35 | 0.38 | 0.10 | 0.13 |

**Table 38:** Power Analysis Results - CBCL Social Problems (Change)

| Method | $n = 250$ | $n = 500$ | $n = 1000$ | $n = 2000$ | a | b | P | $R^2$ |
| --- | --- | --- | --- | --- | --- | --- | --- | --- |
| Network FDR | 0.20 | 0.09 | 0.15 | 0.31 | 99942.11 | 0.47 | 1.94 | 0.38 |
| Network FWER | 0.16 | 0.05 | 0.15 | 0.27 | 99921.56 | 0.58 | 2.64 | 0.49 |
| TFCE | 0.00 | 0.01 | 0.01 | 0.00 | 751.29 | 0.12 | 0.01 | -0.05 |
| Omnibus.cNBS | 0.00 | 0.00 | 0.00 | 0.00 | 751.83 | 2.64 | $2.7 \cdot 10^{-4}$ | -Inf |
| Parametric FDR | 0.00 | 0.00 | 0.00 | 0.00 | 751.30 | 0.16 | 0.01 | -0.14 |
| Parametric FWER | $8.4 \cdot 10^{-4}$ | $8.4 \cdot 10^{-4}$ | $9.8 \cdot 10^{-4}$ | $6.7 \cdot 10^{-4}$ | 751.82 | 2.62 | 0.00 | -22.56 |
| Size | 0.04 | 0.04 | 0.08 | 0.04 | 751.28 | 0.11 | 0.10 | 0.01 |

**Table 39:** Power Analysis Results - CBCL Somatic Complaints (Change)

| Method | $n = 250$ | $n = 500$ | $n = 1000$ | $n = 2000$ | a | b | P | $R^2$ |
| --- | --- | --- | --- | --- | --- | --- | --- | --- |
| Network FDR | 0.13 | 0.18 | 0.29 | 0.07 | 751.29 | 0.10 | 0.34 | -0.02 |
| Network FWER | 0.13 | 0.16 | 0.25 | 0.07 | 751.29 | 0.10 | 0.31 | -0.03 |
| TFCE | 0.00 | 0.00 | 0.01 | 0.01 | 751.59 | 1.54 | 0.01 | 0.92 |
| Omnibus.cNBS | 0.00 | 0.00 | 0.00 | 0.00 | 751.83 | 2.64 | $2.7 \cdot 10^{-4}$ | -Inf |
| Parametric FDR | 0.00 | $8.9 \cdot 10^{-4}$ | 0.00 | 0.00 | 751.56 | 1.39 | 0.00 | 0.80 |
| Parametric FWER | $4.2 \cdot 10^{-4}$ | $2.0 \cdot 10^{-4}$ | $5.0 \cdot 10^{-4}$ | $9.5 \cdot 10^{-4}$ | 751.83 | 2.64 | $9.7 \cdot 10^{-4}$ | 0.45 |
| Size | 0.04 | 0.03 | 0.03 | 0.09 | 751.52 | 1.07 | 0.09 | 0.52 |

**Table 40:** Power Analysis Results - CBCL Withdrawn/Depressed (Change)

| Method | $n = 250$ | $n = 500$ | $n = 1000$ | $n = 2000$ | a | b | P | $R^2$ |
| --- | --- | --- | --- | --- | --- | --- | --- | --- |
| Network FDR | 0.15 | 0.11 | 0.27 | 0.15 | 751.34 | 0.28 | 0.34 | 0.09 |
| Network FWER | 0.11 | 0.11 | 0.22 | 0.15 | 751.37 | 0.44 | 0.29 | 0.30 |
| TFCE | 0.00 | 0.00 | $2.8 \cdot 10^{-4}$ | $6.1 \cdot 10^{-4}$ | 751.82 | 2.61 | 0.00 | -2.67 |
| Omnibus.cNBS | 0.00 | 0.00 | 0.00 | 0.00 | 751.83 | 2.64 | $2.7 \cdot 10^{-4}$ | -Inf |
| Parametric FDR | $5.0 \cdot 10^{-4}$ | 0.00 | $2.5 \cdot 10^{-4}$ | 0.00 | 751.82 | 2.62 | 0.00 | -0.73 |
| Parametric FWER | $2.5 \cdot 10^{-4}$ | $2.8 \cdot 10^{-4}$ | $2.0 \cdot 10^{-4}$ | $5.9 \cdot 10^{-4}$ | 751.83 | 2.63 | $6.4 \cdot 10^{-4}$ | -0.26 |
| Size | 0.01 | 0.01 | 0.01 | 0.02 | 751.41 | 0.74 | 0.03 | 0.38 |

**Table 41:** Power Analysis Results - CBCL Anxious/Depressed (Change)

| Method | $n = 250$ | $n = 500$ | $n = 1000$ | $n = 2000$ | a | b | P | $R^2$ |
| --- | --- | --- | --- | --- | --- | --- | --- | --- |
| Network FDR | 0.13 | 0.16 | 0.31 | 0.11 | 751.30 | 0.15 | 0.36 | 0.02 |
| Network FWER | 0.09 | 0.11 | 0.29 | 0.11 | 751.38 | 0.47 | 0.30 | 0.11 |
| TFCE | 0.06 | 0.09 | 0.02 | 0.01 | 751.28 | 0.10 | 0.09 | -0.09 |
| Omnibus.cNBS | 1.00 | 0.00 | 0.00 | 0.00 | 1.00 | 0.10 | 0.37 | -0.02 |
| Parametric FDR | 0.07 | 0.07 | 0.01 | 0.01 | 751.28 | 0.10 | 0.08 | -0.09 |
| Parametric FWER | 0.00 | 0.00 | $8.1 \cdot 10^{-4}$ | 0.00 | 751.31 | 0.21 | 0.00 | -0.26 |
| Size | 0.07 | 0.11 | 0.12 | 0.11 | 751.36 | 0.40 | 0.20 | 0.60 |

### 6.3 Documentation

Documentation for `PRISME` is available at https://neuroprismlab.github.io/PRISME-Brain-Power-Calculator/. The documentation describes input data format requirements, configuration parameters, and execution workflows, alongside a developer guide detailing the process for implementing new statistical inference methods, overall code architecture, and C++ integration for performance optimization.

### 6.4 Data Preprocessing

Functional images were motion-corrected using SPM5. The data were then iteratively smoothed to an equivalent smoothness of a 2.5 mm Gaussian kernel in order to ensure uniform smoothness across the dataset. White matter and CSF were defined on a MNI-space template brain and eroded in order to minimize inclusion of grey matter in the mask. The template was then warped to subject space using a series of transformations described in the next section. This ensured that mainly grey matter voxels were used in subsequent analyses. The following noise covariates were regressed from the data: linear, quadratic, and cubic drift, a 24-parameter model of motion, mean cerebrospinal fluid signal, mean white matter signal, and mean global signal. Finally, data were temporally smoothed with a zero mean unit variance Gaussian filter (cutoff freq=0.19 Hz). Anatomical data were first skull-stripped using FSL. Functional data for each subject, scanner, and session were linearly registered to the corresponding FLASH images. FLASH images were then linearly registered to MPRAGE images. Next, an average MPRAGE image for each subject was created by linearly registering and averaging all 4 anatomical images (from 2 scanner × 2 sessions) for each subject. These average MPRAGE images were used for an iterative nonlinear registration to MNI space. The use of the average anatomical images and a single nonlinear registration for each subject ensures that any potential anatomical distortions due to the different scanners does not introduce a systematic bias into the registration. The average MPRAGE images were nonlinearly registered to an evolving group average template in MNI space as described previously. All transformation pairs were calculated independently and then combined into a single transform that warps single participant results into common space. From this, all subjects’ images can be transformed into common space using a single transformation, which reduces interpolation error. [NST^+^17]

### 6.5 Computational Environment and Performance Details

`PRISME` was speed benchmarked in a different computing environment to the previous empirical power calculation performed by Noble et al. [NMZS22]. In this section, we provide details about both computing cluster environments.

#### Noble et al. benchmarking

The original implementation benchmarks were conducted on the Farnam High Performance Computing cluster at Yale University via the Simple Linux Utility for Resource Management (SLURM) workload manager. Resources used for calculations were 13 CPUs and 40GB of memory on compute nodes from the general partition (Red Hat Enterprise Linux Server 7.9).

#### PRISME benchmarking

`PRISME` was evaluated on the now-deactivated Discovery cluster at Northeastern University. The Discovery cluster had a total of over 50,000 CPU cores and over 525 GPUs, with hardware consisting of a combination of Intel Xeon (Cascadelake, Skylake, Broadwell, Haswell, Sandybridge, and Ivybridge) and AMD (Zen, Zen2) CPU microarchitectures. Compute nodes are connected via either 10 GbE or high-data-rate InfiniBand (200 Gbps or 100 Gbps). Storage is provided through a high-performance file system with 6 PB of available storage. Resources used for calculations were 25 CPUs and 70 GB of memory. Computational speed comparisons assumed perfect parallelism and used sequential time to account for core differences.

## 7 Theoretical Validation

### 7.1 Model and Assumptions

In this section, we provide formal validation for the power estimation of `PRISME` and show convergence to the true statistical power. We present the key assumptions, proof strategy, and proof elements in accessible terms before the formal mathematical treatment. The explanations are more detailed due to the potential lack of familiarity of the neuroscience audience with measure theory and the mathematical concepts used to validate `PRISME`.

#### Lebesgue Measurement

The Lebesgue measure is the volume created by a subset of real numbers. One can think of the Lebesgue measure as a replacement for the differential element in integration when calculating a volume. For example, if we consider the real line with *x*_1_, *x*_2_ ∈ ℝ, *x*_2_ > *x*_1_, then this is true for the Lebesgue measure:

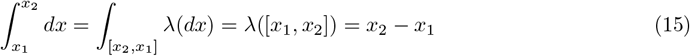

Similarly to the differential element *dx*, the Lebesgue measure is zero for points. For higher dimensions, for example, when integrating over areas in ℝ ^2^, the Lebesgue measure of lines and dots is zero, because their area is zero.

For the validation, we need the Lebesgue measure to characterize two things: (1) which types of probability distributions from population measurements can be applied `PRISME` to produce valid results, and (2) which types of statistical tests produce valid convergence. The theoretical guarantee provided by `PRISME` holds for continuous probability distributions with respect to the Lebesgue measure and for statistical tests with well-behaved decision boundaries (boundaries with zero Lebesgue measure). These conditions ensure convergence by repeated subsampling.

#### Distribution Assumption

We model individual subject measurements as draws from a distribution *F* in ℝ ^*V*^, where *V* is the number of variables, that is absolutely continuous with respect to the Lebesgue measure. Intuitively, this means the probability of observing any exact value is zero on a multivariate generalization of the fact that continuous random variables assign zero probability to single points. In the mathematical proof, this is written as:

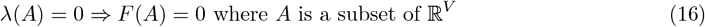

As an example, take the one-dimensional normal distribution in ℝ. In the real line, the Lebesgue measure of points is zero, and the normal distribution has probability zero of taking an exact value with infinite precision.

This assumption is true for all distributions commonly used in neuroimaging, including the normal, Student’s t, chi-squared, and exponential distributions, and excludes only discrete distributions with fixed probability masses at specific values.

#### Independence Between Subjects Assumption

We assume that subsampled subjects are independent and identically distributed draws from the empirical distribution *F*_*N*_ . The empirical distribution *F*_*N*_ is the distribution obtained from the *N* subjects drawn from the entire population to construct the large dataset used by `PRISME`.

In practice, `PRISME` subsamples without replacement, which introduces weak dependence, as the following draws become dependent on the subjects removed by previous draws. However, when the dataset size *N* is much larger than the subsample size *n*, this dependence becomes negligible, and the i.i.d. approximation is justified. The proof could likely be extended to include this dependence, but this assumption greatly simplifies the analysis. A more detailed analysis without this assumption is more in the scope of a completely dedicated theoretical work of resampling procedures for power calculation.

#### Statistical Test Assumption

We model statistical tests as deterministic measurable functions that receive all the *n* subject values from a subsampled repetition. When modelled like this, the statistical tests define regions across the space of possible values of subjects where the test rejects the null hypothesis and where it does not. The notation of the region of all the subsampled subject values is 𝒟 _*v*_. This specific type of modelling statistical tests is not novel, and it is well established in statistical theory by Neyman and Pearson [NP33].

Although parametric tests are deterministic, permutation tests are normally not. They become deterministic if all possible permutations of the data are used to compute the null. However, in practical and empirical terms, it is only necessary to compute enough permutations that the test produces consistent results across equal subject measurements.

We assume the boundary between rejection and non-rejection is a thin surface rather than a thick region, and that the decision region is not scattered into isolated points. As a 2-dimensional example, imagine a valid boundary as a line or collection of lines, not necessarily straight or finite, separating the regions of rejection and non-rejection. Formally, this is stated as the Lebesgue measure of the boundary being zero. This assumption holds for all statistical tests currently used in neuroscience and is included only to guard against non-empirical counterexamples. In the mathematical proof, this is written as:

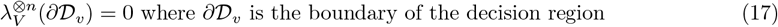

Here, the Lebesgue measure 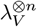 is the Lebesgue measure of a volume in (ℝ ^*V*^)^*n*^. The space of measurements for a sub sampled experiment is (ℝ ^*V*^)^*n*^ which represents the measurements from *n* subjects in the subsample.

#### 7.1.1 Proof Strategy

The proof proceeds in three steps:

##### Step 1: Power as an Expected Value

Firstly, we show that power can be written as the expected number of times the subject measurements fall in the region of rejection of the test. This uses an established relationship between probabilities and the expected value of an indicator function.

##### Step 2: Convergence of the product distribution

Since subjects are independent, the joint distribution of an experiment with *n* subjects is the product distribution *F* ^⊗*n*^. Similarly, experiments using the empirical distribution follow 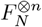. The product distribution is the distribution one gets when modelling multiple draws from multiple random variables. For example, the probability distribution of getting multiple flips from a coin is the product distribution of each individual coin flip.

In the mathematical proof, this step is saying that because this statement is true:

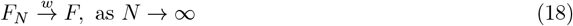

The following statement is implied:

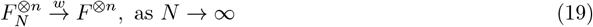

This essentially means: Because the distribution of each subject drawn from the dataset converges to the distribution of the real population as the size of the dataset increases, the distribution of an experiment with *n* subjects converges to the distribution of an experiment with the real distribution as the size of the dataset increases.

Finally, we apply the Portmanteau theorem [VdV00] to show that expectations under the empirical product distribution 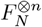 converge to expectations under the true product distribution *F* ^⊗*n*^ as *N → ∞*. The well-behaved boundary assumption assures that the theorem can be applied.

##### Step 3: Sample Average Converges to Expectation

We use the law of large numbers to show the proportion of successful detections across *R* repetitions converges to the expected value of detections as *R* → ∞.

#### 7.1.2 Scope

The proof establishes that `PRISME`’s power estimates converge to true power as the dataset size *N* and number of repetitions *R* increase. The definition of convergence shows that there exists a **finite** *N* for which the empirical error becomes negligible. This establishes `PRISME` as a viable power calculator, given enough subjects in the large dataset. Unfortunately, we do not analyse exactly which value of *N* might be enough to result in negligible empirical errors. This would require either convergence proofs and analyses that are out of the scope of this work.

The proof applies to variables where the null hypothesis is false (true effects exist), as statistical power is only defined in this case. Although not directly stated, it can be inferred from the proof that for variables with no true effect, the detection rate estimated by `PRISME` converges to the proportion of errors that would be obtained by applying the statistical test to a variable with no true effect (with respect to the used null hypothesis).

### 7.2 Theorem

The main validation theorem follows:

#### Theorem 1

(Convergence of PRISME Power Estimation). *Let F be a probability distribution on ℝ*^*V*^, *V* ∈ *ℕ*_*>*0_, *that is absolutely continuous with respect to the Lebesgue measure. Let F*_*N*_ *denote the empirical distribution estimated from N* ∈ *ℕ*_*>*0_ *i*.*i*.*d. realisations of F* .

*Let n* ∈ *ℕ*_*>*0_. *Let* 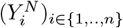 *be a tuple of n i*.*i*.*d. random vectors drawn from F*_*N*_ *in* (ℝ^*V*^)^*n*^. *For* 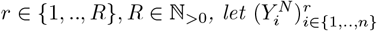 *denote the r-th independent realization of this tuple*.

*Let v be a variable index for which the null hypothesis is false. Let* 𝒮_*v*_ : (ℝ^*V*^)^*n*^ *→* 0, 1 *be a well-behaved statistical test*.

*Then, almost surely:*

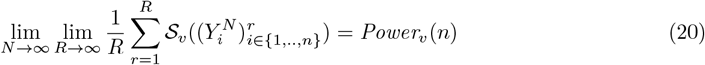

*where Power*_*v*_(*n*) *is the probability of correctly rejecting the null hypothesis at the v-th variable when it is false, given a sample size of n*.

The result follows from Lemmas 3, 4, and 5. By Lemma 5, the sample average converges to the expectation under *F*_*N*_ as *R* → ∞. By Lemma 4, this expectation converges to the expectation under *F* as *N* → ∞. By Lemma 3, this expectation equals the true statistical power.

### 7.3 Proof

Let *n* ∈ ℕ_*>*0_. Let *S* = {1, 2, …, *n*} be a set of indices.

Let 𝒱= {1, 2, …, *V*} denote a finite set of indexes for the variables and *V* ∈ *ℕ*_*>*0_ denote the number of variables.

Let (*X*_*i*_)_*i*∈*S*_ be a tuple of independent and identically distributed (i.i.d.) random vectors in ℝ^*V*^ . Each *X*_*i*_ follows the distribution *F* :

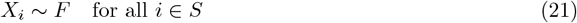

such that the distribution *F* is absolutely continuous with respect to the Lebesgue measure:

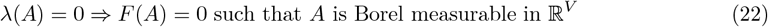

Each random vector *X*_*i*_ can be written as:

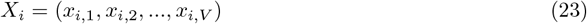

where *x*_*i,j*_ ∈ ℝ denotes the measurement of variable *j* for subject i.

Let *N* ∈ *ℕ*_*>*0_.

Let 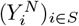 be a tuple of random vectors in ℝ^*V*^ . Each 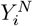 follows an empirical distribution estimated from the distribution of *X*_*i*_ using *N* realizations. Consequently, from the definition of *X*_*i*_, one can conclude that the elements of the tuple 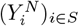 are also i.i.d. and they follow the distribution *F*_*N*_ where *F*_*N*_ is the empirical distribution estimated with *N* realisations:

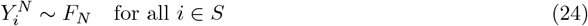

We assume proper sampling is performed, which means that *F*_*N*_ is fixed on an outcome of the probability one set that converges weakly to *F* :

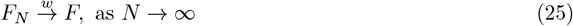

This assumption is satisfied almost surely when *F*_*N*_ is constructed as the empirical distribution from random sampling of *F* [Bil13].

Let *H* be a null hypothesis. Let 𝒮_*v*_ : (ℝ^*V*^)^*n*^ → {0, 1} be a Borel measurable function that tests for null hypothesis *H* that returns 1 if the variable *v* across all (*X*_*i*_)_*i*∈*S*_ was declared significant by the method (the null hypothesis is false at variable index *v*).

The function 𝒮_*v*_ is deterministic. Therefore, it is only well defined for parametric methods and non-parametric permutation methods that use all possible permutations to estimate the null. For empirical purposes, it is necessary to have enough permutations that the non-parametric method returns the same value across multiple executions for the same measurements.

Let 𝒟_*v*_ ⊆ (ℝ^*V*^)^*n*^ be the set of subject measurements for which for all *e* ∈ 𝒟_*v*_

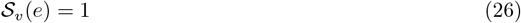

We call the set 𝒟_*v*_ the decision region for variable index *v*. We assume throughout that 𝒮_*v*_ produces a well-behaved decision region. This assumption is satisfied by all standard statistical inference methods. Violations arise only from pathological constructions with no practical application. The definition of a well-behaved statistical test follows:

#### Defintion 2

(Well-behaved statistical test). *A deterministic statistical test* 𝒮_*v*_ : (ℝ^*V*^)^*n*^ 0, 1 *is called well-behaved if the following conditions hold:*

*1. 𝒮*_*v*_ *is Borel measurable*

*2*. 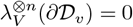, *where* 𝒟_*v*_ = *{e* ∈ (ℝ^*V*^)^*n*^ : 𝒮_*v*_(*e*) = 1}

If 𝒮_*v*_ is Borel measurable, the decision region 𝒟_*v*_ is a Borel measurable set, and consequently its boundary ∂ 𝒟_*v*_ is also Borel measurable. Therefore, the boundary condition in the definition is well-defined.

Without loss of generality, consider variables with positive true effects in the population. Let 𝒱^*H*^ denote the set of variable indexes for which the null hypothesis *H* is false (rejected by the null).

Statistical power is defined in the standard way as the probability of correctly rejecting the null hypothesis when it is false. For a variable index *v* ∈ 𝒱^*H*^ :

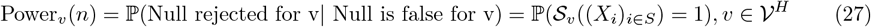

Now that our model is defined, we prove the first lemma. `PRISME` calculates power in terms of expectations by resampling. So we start by showing that the power of a method can be transformed into an expectation.

#### Lemma 3.

*For any variable index v* ∈ 𝒱, *we have:*

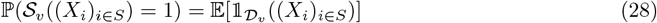

*Proof*. From the definition of 𝒟_*v*_ (Equation (26)), one has that 𝒮_*v*_((*X*_*i*_)_*i*∈*S*_) = 1 if and only if (*X*_*i*_)_*i*∈*S*_ ∈ 𝒟_*v*_. Finally, from a fundamental property of expected value and indicator functions, we have:

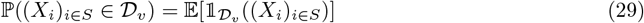

and the proof is concluded

Now, to continue the proof, we prove that the random variables that follow empirical distributions *F*_*N*_ converge to *F* as the sample size increases to infinity.

#### Lemma 4.

*It is:*

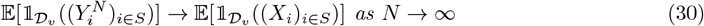

*Proof*. Note 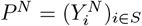 and *P* = (*X*_*i*_)_*i*∈*S*_.

As the random vectors in 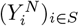 are i.i.d, the probability distribution of *P*^*N*^ is the product distribution of *F*_*N*_ :

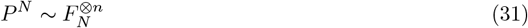

Equally for *P* :

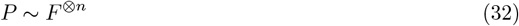

From the sampling assumption and the definition of *F*_*N*_, we have the weak convergence of the empirical distribution [Bil13]:

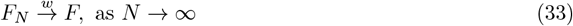

Consequently, as weak convergence is preserved over finite products [Bil13], we have The following:

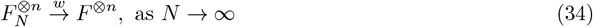

From the absolute continuity of *F*, for any Borel measurable set *A*_*V*_ in *R*^*V*^, the measure *λ*_*V*_ (*A*) = 0 implies *F* (*A*) = 0. Therefore, from the preservation of the absolute continuity for the product measures [Bil13], the following statement holds:

For any set Borel measurable set *A*_*V n*_ in (*R*^*V*^)^*n*^:

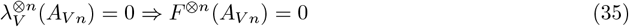

From the definition 𝒟_*v*_ in Definition 2, it follows for the Lebesgue measure of ∂𝒟_*v*_:

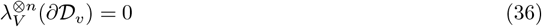

Therefore, from Equation (35), it implies for the measure *F* ^⊗*n*^:

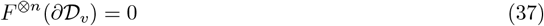

Furthermore, the indicator function 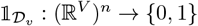 takes value 1 for points inside 𝒟_*v*_ and value 0 for points outside 𝒟_*v*_. Consequently, 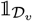 is continuous everywhere except at the boundary ∂ 𝒟_*v*_, where the function value changes discontinuously from 0 to 1.

Therefore, due to the three following statements:

- *P*^*N*^, *P* are in (ℝ^*V*^)^*n*^
- From Equation (34), 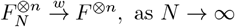
- From Equation (37), *F* ^⊗*n*^(∂𝒟_*v*_) = 0, which means 𝒟_*v*_ is a continuity set with respect to *F* ^⊗*n*^. One can invoke the Portmanteau theorem [Kle08] to conclude that the following convergence holds:

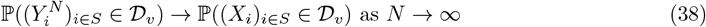

Consequently, from a fundamental property of the expected and the indicator functions, one has:

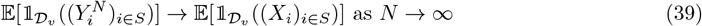

Finally, we conclude the theorem proof with a final lemma by using the law of large numbers.

#### Lemma 5.

*Let* ℛ = {1, 2, 3, 4, 5, *· · ·, R}. Let* 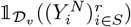 *denote an independent draw from the random variable* 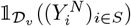.

*It is:*

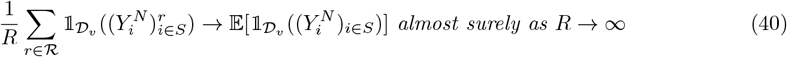

*Proof*. Since for all *r* ∈ ℛ, one has that 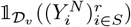 is a draw from the independent random variable 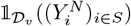 by the law of large numbers:

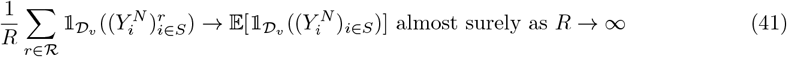

With Lemmas 4, 5, and 3, the proof is then concluded, as for *v* ∈ 𝒱^*H*^, one has:

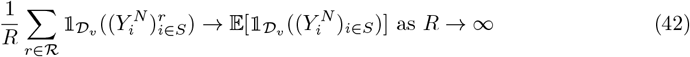

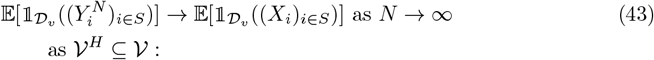

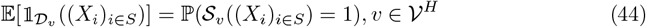

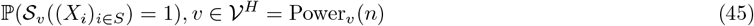

and the proof is concluded.

